# Cyclin Y Overexpression Drives a Fatal Metabolic Syndrome via Defective Glucose Homeostasis

**DOI:** 10.64898/2025.12.15.694124

**Authors:** Luis R. López-Sainz, Estefanía Ayala-Jimenez, Agustín Sánchez-Belmonte, Sagrario Ortega, Eduardo Caleiras, Eduardo Zarzuela, Marta Isasa, Clara M. Santiveri, Ramón Campos-Olivas, Guillermo de Cárcer, Marcos Malumbres

**Affiliations:** Cell Division and Cancer group, Spanish National Cancer Research Center (CNIO), Melchor Fernández Almagro 3, Madrid 28029, Spain; Cancer Cell Cycle group, Vall d’Hebron Institute of Oncology (VHIO), Natzaret 115-117, Barcelona 08035, Spain; Centro de Investigación Biomédica en Red de Cáncer (CIBERONC), Instituto de Salud Carlos III, Spain; Mouse Gene Editing Unit, Spanish National Cancer Research Center (CNIO), Melchor Fernández Almagro 3, Madrid 28029, Spain; Histopathology Unit. Spanish National Cancer Research Centre (CNIO), Madrid, Spain; ProteoRed-ISCIII, Unidad de Proteómica, Centro Nacional de Investigaciones Oncológicas (CNIO), Melchor Fernández Almagro 3, 28029, Madrid, Spain; Spectroscopy and Nuclear Magnetic Resonance Unit, Structural Biology Programme, Centro Nacional Investigaciones Oncológicas (CNIO), Madrid, Spain; Cell Cycle and Cancer Biomarkers Group, Department of Cancer Biology, Instituto de Investigaciones Biomédicas Sols-Morreale (IIBM, CSIC-UAM), Arturo Duperier 4, 28029, Madrid, Spain. Conexion-Cancer Hub (CSIC); Translational Cancer Research Group (Area 3-Cancer), Instituto Ramón y Cajal de Investigación Sanitaria (IRYCIS), Ctra. de Colmenar Viejo, Km.9, 28034, Madrid, Spain; ICREA, Passeig Lluís Companys 23, Barcelona 08010, Spain

**Keywords:** cyclin Y, cyclin-dependent kinase, glucose metabolism, gluconeogenesis, liver damage

## Abstract

Cyclin Y (CCNY) is a membrane-associated, non-canonical cyclin best known for regulating WNT/beta-catenin signaling via the recruitment of CDK14/16 protein kinases. Whereas its role in the activation of members of the atypical CDK14-18 subfamily is established, its function in systemic metabolism remains poorly defined. Here, we report that CCNY overexpression drives a fatal metabolic syndrome. Using a novel inducible knock-in mouse model, we demonstrate that CCNY overexpression causes severe cachexia and profound hypoglycemia, resulting in death of adult mice independent of tumor burden. While these mutant mice maintain normal food intake and hepatic synthetic function, they succumb to metabolic starving state. Proteomics reveals that CCNY interacts with and stabilizes Pyruvate Dehydrogenase Kinase 4 (PDK4), in agreement with defective pyruvate use at mitochondria and enforcing a Warburg-like shift to aerobic glycolysis. Phosphoproteomic analysis indicates activation of apoptotic pathways and defective phosphorylation of several enzymes critical for glycolysis and gluconeogenesis as well as amino acid metabolism in addition to other metabolic routes. Altogether, these data suggest that CCNY overexpression uncouples nutrient sensing from utilization, a finding with possible therapeutic implications in CCNY-high cancers with similar changes in metabolic pathways.

## INTRODUCTION

Cyclins are a large family of approximately 30 proteins characterized by a particular amino acid sequence called the cyclin box, which is necessary for the enzymatic activity of cyclin-dependent kinases (CDKs)(Malumbres, 2014; Martinez-Alonso & Malumbres, 2020; Quandt *et al*, 2020). Cyclin Y (CCNY) and CCNY-like 1 CCNYL1) represent a particular group characterized by the presence of a single cyclin fold, while most other cyclins have two (Jiang *et al*, 2009; Mikolcevic *et al*, 2012a). Y-type cyclins are also unique for the presence of myristoylation and palmitoylation signals, placed in the N-terminal domain, which are essential for cyclin Y translocation to the plasma membrane (Malumbres, 2014; Seo *et al*, 2022). Two additional CCNY-like sequences are present in humans, although they are typically characterized as pseudogenes (CCNYL2 and CCNYL3; (Davidson *et al*, 2009)).

Y-type cyclins are known to bind and activate CDK14 and CDK16 (Davidson *et al*., 2009; Mikolcevic *et al*, 2012b; Shehata *et al*, 2012). CCNY, CCNYL1 and their CDK partners act upstream of WNT, and they can regulate the canonical WNT/β-catenin signaling pathway and WNT-mediated stabilization of proteins (WNT/STOP) (Acebron *et al*, 2014; Davidson & Niehrs, 2010; Zeng *et al*, 2016). This β-catenin-independent pathway has been described to be essential for neurogenesis in the neocortex during development (Da Silva *et al*, 2021), and cyclin Y is known to be involved in synapsis and neuronal homeostasis (Seo *et al*., 2022). However, lack of the murine *Ccny* gene mostly results in metabolic defects, characterized by altered lipid metabolism, reduced body weight and fat content, defective adipocytes differentiation and inhibition of liver insulin signaling (An *et al*, 2015). Concomitant ablation of *Ccny* and *Ccnyl1* results in mouse embryonic lethality at E16.5 as a consequence of reduced WNT signaling and defective proliferation and regeneration of stem cells in several organs (Da Silva *et al*., 2021; Zeng *et al*., 2016), suggesting redundant functions of CCNY and CCNYL1 in WNT signaling.

CCNY and CDK16 overexpression has been reported in different cancer types (Liu *et al*, 2016; Liu *et al*, 2025; Shi *et al*, 2018; Sun *et al*, 2014; Yan *et al*, 2016; Yue *et al*, 2011; Zhao *et al*, 2021), although it is currently unclear whether these observations simply correlate with cell cycle activation, or these proteins display functional oncogenic activities. In this report, we generated an inducible cyclin Y-overexpressing mouse model by introducing a single copy of a CCNY cDNA under a Tet-responsive element in the mouse genome. Unexpectedly, overexpression of CCNY is lethal in adult mice with complete penetrance in a dose-dependent manner. This phenomenon is not related to the presence of tumors but rather to metabolic alterations that dysregulate glucose metabolism leading to liver damage and death by metabolic starvation in adult mice.

## Results

### Premature death and metabolic alterations in CCNY-overexpressing mice

We generated a doxycycline (Dox)-inducible, whole-body *CCNY* knock-in mouse model by placing a cassette containing the human *CCNY*-FLAG cDNA under the control of a tetracycline-responsive element (TRE) in the mouse *Col1A1* locus [*Col1A1*(Y) allele; Fig. 1a and Suppl. Fig. 1a]. The *CCNY* gene was fused to FLAG in the C-terminal sequence to avoid the alteration of the myristoylation and palmitoylation signal located at the N-terminal, essential for cyclin Y localization and function (Seo *et al*., 2022). The Dox-dependent rtTA operator was expressed in the ubiquitously expressed *Rosa26* locus [*Rosa26*(rtTA) allele; Fig. 1a] following a strategy reported previously (Beard *et al*, 2006). Overexpression of CCNY-FLAG was evident in vivo as early as 12 hours after initiation of Dox diet in multiple tissues by using CCNY- or FLAG-specific antibodies (Fig. 1b). CCNY was highly overexpressed after 6 days of Dox diet in most tissues except from brain, and heart (Suppl. Fig. 1b), a limitation previously reported for this knock-in strategy (Beard *et al*., 2006). CCNY was mostly defected in cellular membranes suggesting that CCNY localization is well preserved in CCNY-FLAG-overexpression mice (Fig. 1c).

**Fig. 1.**
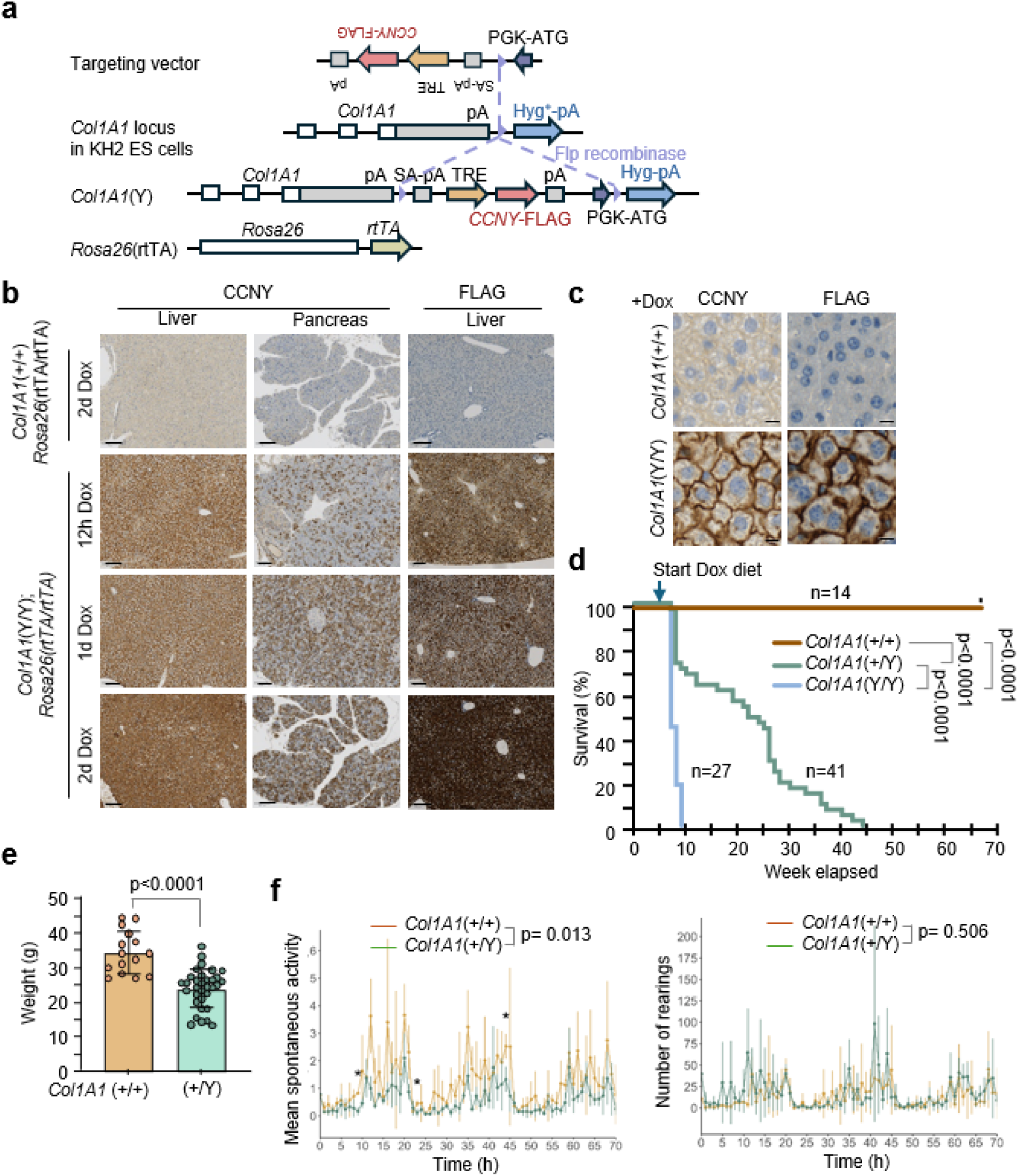
Cyclin Y over-expression reduces body weight and compromises survival. **a)** Illustration of cyclin Y knock-in mouse model with the *Rosa26* locus located at chromosome 6 and the *Col1A1* gene locus, containing the h*CCNY-FLAG* cDNA sequence, placed at chromosome 11. **b)** Immunological detection of CCNY and FLAG in liver and pancreas of mice at the indicated time points of doxycycline (Dox) diet administration. Scale bar, 100 μm. **c)** Immunohistochemistry using CCNY- and FLAG-specific antibodies in the liver 24 h after Dox diet administration. Scale bar, 10 μm. **d)** Survival curves of *Col1A1*(+/+) (n=14), *Col1A1*(+/Y) (n=41) and *Col1A1*(Y/Y) (n=27) mice. All animals contain the *Rosa26*(rtTA/rtTA) alleles (not indicated from now on). Survival analysis was performed using the Mantel-Cox test for the compared conditions. **e)** Body weight of *Col1A1*(+/Y) mice and control littermates at humane endpoint (n = 15-33 per genotype). Each dot represents a single mouse. Data are expressed as means ± SD (two-tailed unpaired Student’s t-test). **f)** Time course analysis of spontaneous activity and rearing activity in control and *Col1A1*(+/Y) mice using metabolic cages. Mice were on Dox diet for 100 days (n = 3 per genotype). Each time point was analyzed using two-tailed unpaired t-test and mixed-effect analysis with Šidák’s multiple comparison test for condition effect analysis (*p<0.05).

Unexpectedly, the median survival of *Rosa26*(rtTA/rtTA); *Col1A1*(Y/Y) mice [from now on *Col1A1*(Y/Y) mice] was only 7 weeks after 1 week of induction, meanwhile it was 22 weeks (16 weeks of induction) in *Col1A1*(+/Y) mice, confirming a dose-dependent response (Fig. 1d). At humane endpoint, weight was significantly reduced in *Col1A1*(Y/Y) mice (Suppl. Fig. 1c), and this observation correlated with reduced food consumption after two days on Dox diet, whereas no reduction was observed in *Col1A1*(+/+) mice under the same diet (Suppl. Fig. 1d). In open field experiments, *Col1A1*(Y/Y) mice displayed reduced mobility (Suppl. Fig. 1e), without alterations in motor activity (Suppl. Fig. 1f). We did not observe structural defects in the gut, brain or peripheral nervous system (Suppl. Fig. 1g), and whether these alterations are cause or consequence of metabolic or neuronal defects is unclear.

The reduction in body weight was also obvious in *Col1A1*(+/Y) heterozygous mutants at humane endpoint (Fig. 1e). However, while we observed some variability in food uptake during the first days on Dox diet, no differences were observed after the first week of treatment (Suppl. Fig. 2a), indicating that death in these CCNY-overexpressing mice was not due to reduced food consumption. We also used metabolic cages for a deeper analysis of alterations in *Col1A1*(+/Y) mice 100 days after CCNY induction. No differences were observed in food intake or water consumption between heterozygous and control mice (Suppl. Fig. 2b). Similarly, when corrected by weight, *Col1A1*(+/Y) and control mice had similar respiratory quotient and energy exchange (Suppl. Fig. 2c). However, we did observe less activity in *Col1A1*(+/Y) mice, with no differences in rearing count (Fig. 1f). Finally, no differences in the number of white blood cells (WBC), red blood cells (RBC) and the hematocrit (HCT) were observed in heterozygous *Col1A1*(+/Y) mice. However, we detected a reduction of hemoglobin (HGB), mean corpuscular volume (MCV) and mean corpuscular hemoglobin (MCH) (Suppl. Fig 2d), suggesting a reduction of circulating hemoglobin and the size of red blood cells, without affecting respiratory quotient (Suppl. Fig 2c).

Histological characterization of *Col1A1*(+/Y) mice at humane endpoint suggested a nearly complete replacement of exocrine pancreas by fatty tissue (Fig. 2a). Additionally, these mutant mice presented increased intestine length (Fig. 2b), a possible consequence emerging from increased food consumption, lower absorption rate, viral infection, short bowel syndrome or some metabolic conditions (Barron *et al*, 2018; Hou *et al*, 2024; Stojanović *et al*, 2021). We also observed accumulation of iron, a marker of ferroptosis, in mutant livers (Fig. 2c). Analysis of blood markers indicated no significant differences in albumin, urea nitrogen or alkaline phosphatase levels (Suppl. Fig. 2e), but significantly increased alanine aminotransferase (ALT, which converts alanine into pyruvate) indicating liver damage. Both gamma-glutamyl transferase and bile acids (synthetized from cholesterol) were significantly increased, a hallmark of cholestasis (Fig. 2d). The upregulation of bile acids was not accompanied by increased cholesterol levels in blood. Instead, *Col1A1*(+/Y) mice presented reduced levels of cholesterol and triglycerides (TG; Fig. 2e), suggesting liver damage was accompanied by reduction of lipid generation at the liver or lipid circulation in blood.

**Fig. 2.**
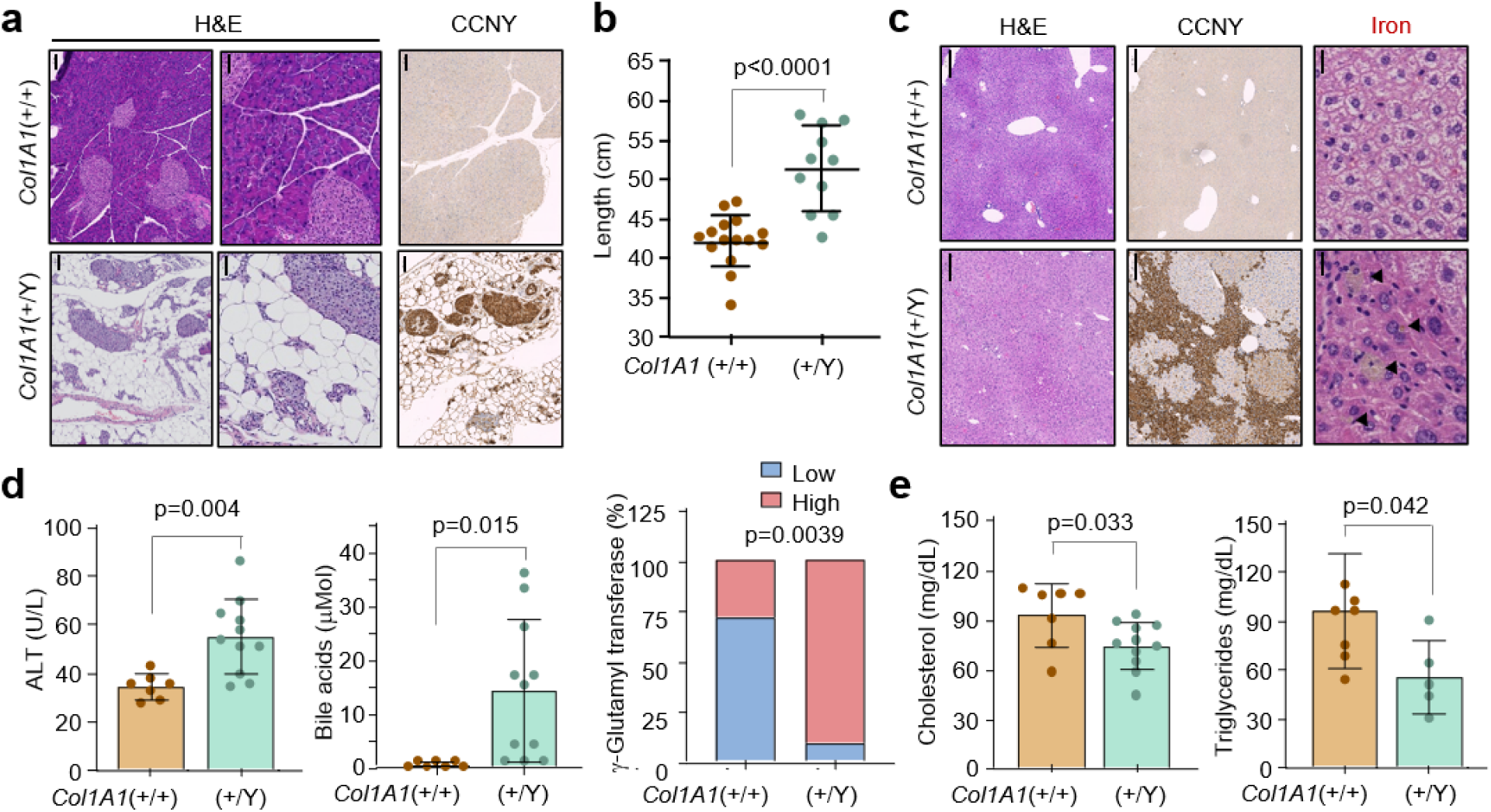
Morphological and physiological alterations in CCNY-overexpressing mice. **a)** Histological images of H&E and CCNY immunodetection in pancreatic samples from *Col1A1*(+/Y) mice at humane endpoint, and paired control samples. Scale bar, 100 μm (left and right panels); 50 μm (middle panels). **b)** Intestine length (in cm) in *Col1A1*(+/Y) mice at humane endpoint and paired control samples (n=11-15 mice per genotype; Two-tailed unpaired Student’s t-test). **c)** Representative histological images of H&E (left) and immunodetection of CCNY (middle) in livers from *Col1A1*(+/Y) mice at humane endpoint and paired control samples. The right panel shows iron accumulation in similar sections. Scale bars, 200 μm (left, middle) and 20 μm (right). **d)** Bloodstream levels of the indicated markers in *Col1A1*(+/Y) mice at humane endpoint and paired control samples. **e)** Bloodstream levels of cholesterol and triglycerides in similar samples. In d,e, each dot represents a single mouse (n=5-11 mice per genotype). In d (γ-glutamyl transferase), “Low” represents 0 U/L and “High” 5 U/L (n = 7-11 per genotype). Data are expressed as means ± SD (two-tailed unpaired Student’s t-test).

### CCNY overexpression results in altered glucose homeostasis

At humane endpoint, CCNY-overexpressing mice displayed a significant reduction in blood glucose level (Fig. 3a). Time course analysis revealed that reduction of blood glucose was an early phenotype that could be observed as early as day 4 after induction of CCNY expression, and that increased with time (Fig. 3b). In these mutant mice, insulin was slightly reduced 6 days after CCNY overexpression and this reduction sustained through life (Fig. 3c). In addition, glucagon was greatly elevated at short and long time points (Fig. 3c), suggesting a consistent physiological response to reduced glucose levels by α-cells in pancreatic Langerhans islets. *Col1A1*(+/Y) mice displayed reduced glucose accumulation after a glucose tolerance test (GTT; Fig. 3d), and after a complementary test in which fasted mice were injected intraperitoneally with sodium pyruvate (Pyruvate tolerance test, PTT; Fig. 3e) suggesting hepatic glucagon resistance and defects in liver gluconeogenesis despite the systemic starvation signal (high glucagon).

**Fig. 3.**
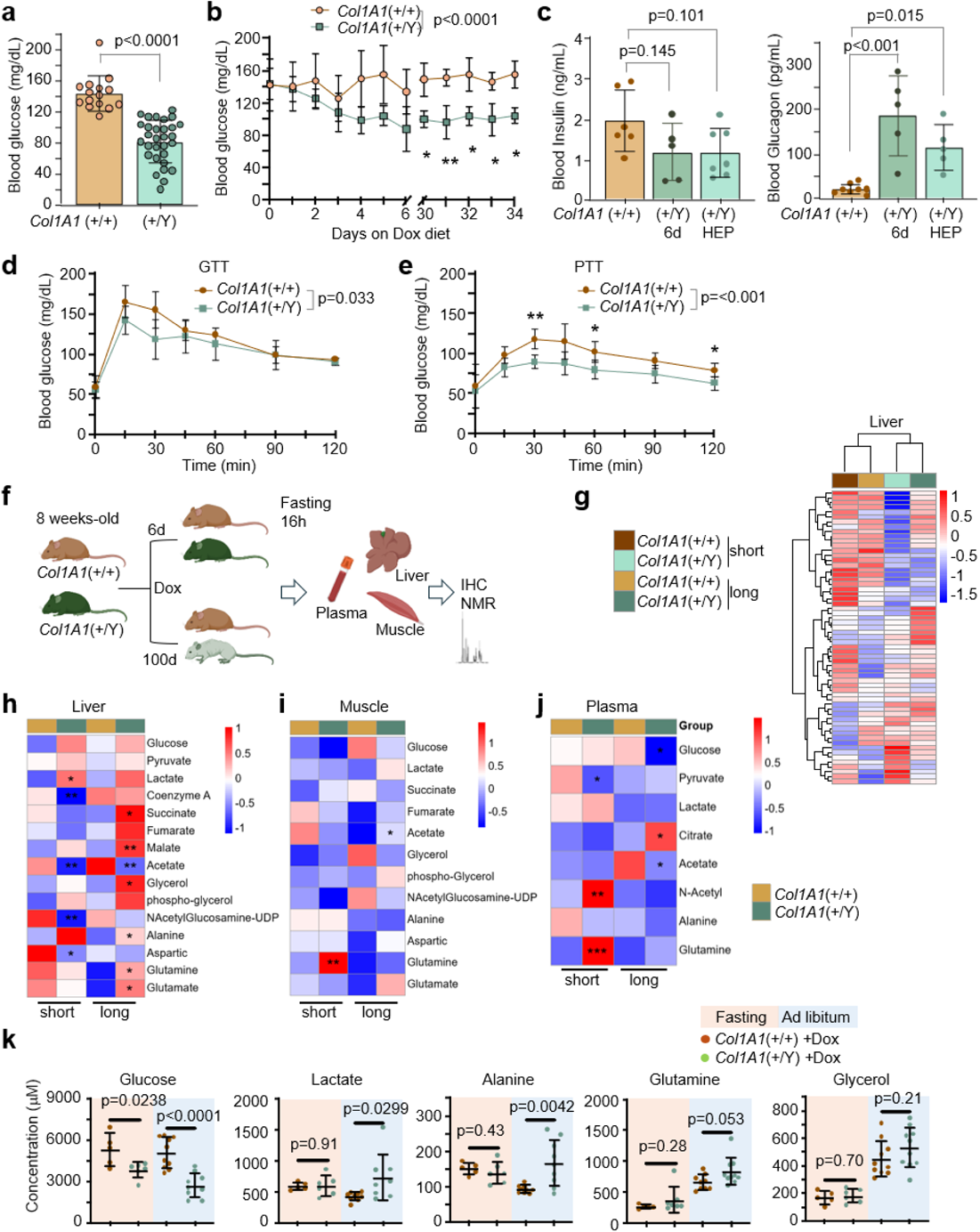
Altered glucose homeostasis and regulation in CCNY overexpressing mice. **a)** Blood glucose levels of *Col1A1*(+/Y) mice at humane endpoint and in paired controls. Each dot represents a single mouse (n=15-30 mice per genotype). Data are expressed as means ± SD (two-tailed unpaired Student’s t-test). **b)** Time course of blood glucose levels following Dox diet administration (n=5-8 mice per genotype). Data were analyzed using mixed-effect analysis with Šidák’s multiple comparison test (*p<0.05; **p<0.005). **c)** Blood levels of insulin and glucagon in control mice and *Col1A1*(+/Y) mice after 6 days on Dox diet and at humane endpoint (HEP). Each dot represents a mouse (n=4-8 mice per group; two-tailed unpaired Student’s t-test;). **d)** Glucose Tolerance Test (GTT). Time course of blood glucose levels of fasted mice injected with glucose (n=8 mice per genotype). **e)** Pyruvate Tolerance Test (PTT). Mice were treated with sodium pyruvate as in the previous assay (n=8 mice per genotype). In d,e, data were analyzed by mixed-effect analysis with Šidák’s multiple comparison test. **f)** Schematic representation of NMR assays, including annotated experimental groups and collected tissue. **g**) Unsupervised clustering of metabolites analyzed by NMR in CCNY-overexpressing and control livers after short or long treatments. **h-j)** Heatmap of z-score corrected data from NMR metabolite concentrations measured in liver (**h**), muscle (**i**) and plasma (**j**). In h-j, groups and time points are indicated (n=5-6 mice per group), and data were analyzed using two-tailed unpaired Student’s t-test of mutant vs. control mice after short or long exposure to Dox (*p<0.05; **p<0.005; ***p<0.0005). **k)** Approximate concentrations of the indicated metabolites in plasma from fasted or ad libitum-fed *Col1A1*(+/Y) or control mice after long Dox exposure. Each dot represents a single mouse (n=5-10 mice per genotype). Data is expressed as means ± SD, and statistical analysis was performed using two-tailed unpaired Student’s t-test.

To analyze more in detail the dynamic response to CCNY overexpression, we subjected control and heterozygous mutant mice to Dox diet for 6 days (short) or 100 days (long). We then fasted mice for 16 hours to synchronize metabolism and avoid variation due to food consumption. We collected liver, muscle and plasma, and analyzed these samples using nuclear magnetic resonance spectroscopy (NMR; Fig. 3f). CCNY was overexpressed at early time points in the liver but not in muscle (Suppl. Fig. 3a). After long exposure to Dox, CCNY was overexpressed both liver and muscle, although livers contained patches of CCNY-positive and CCNY-negative cells suggesting selection against overexpression of this protein (Suppl. Fig. 3a). Clustering analysis of NMR data in the liver suggested specific metabolites altered in CCNY-overexpressing mice that were maintained through life (Fig. 3g). NMR data in liver extracts suggested increase of intracellular lactate and alteration of acetate after short treatment (Fig. 3h). After long treatment, glycerol (a molecule between lipid and glucose metabolism), and molecules connected to intermediates of the citric acid (TCA) cycle, alanine (which can be converted to pyruvate by ALT), glutamine and glutamate were also altered (Fig. 3h). These differences were less obvious in samples from muscle (Fig. 3i), whereas glucose and acetate were significantly downregulated in the plasma after long exposure (Fig. 3j). NMR analysis provided evidence for additional alterations in nucleotide synthesis and amino acids levels (Suppl. Fig. 3b,c). Moreover, lipid metabolism was altered in the liver, showing an increment of polyunsaturated fatty acid (PUFA), diacylglycerol (DAG), inositol lipids and cholesterol free (used for the synthesis of bile acids), and a reduction of circulating lipids, confirming previous results (Suppl. Fig. 3d). As expected from the previous observations, *Col1A1*(+/Y) mice showed a more pronounced defect in plasma glucose levels in ad libitum conditions. We also found higher bloodstream levels of lactate, alanine and glutamine after long treatment in ad libitum, but not fasting conditions (Fig. 3k). These molecules share the capability of being transformed into glucose in the liver, suggesting, all together, defects in gluconeogenesis and a preference for glycolysis in CCNY-overexpressing livers.

For a deeper cell-autonomous biochemical analysis of glucose metabolism, we generated cultured embryonic fibroblasts (MEFs) from *Col1A1*(Y/Y) and control mice. Overexpression of CCNY in these cells was obvious as soon as 6 h on 1 μg/mL Dox (Fig. 4a). GLUT1, the main glucose transporter in MEFs, was upregulated upon CCNY induction (Fig. 4b), suggesting that CCNY-overexpressing cells may be exceedingly up taking glucose from the bloodstream. MEFs were then maintained during 48h in media containing ^13^C-Glucose and cell extracts at the final endpoint were analyzed to observe the variation of cell media and intracellular metabolites. Induction of CCNY led to increase in intracellular glucose, as well as glycerol, glutamine, and several TCA metabolites (Fig. 4c). However, glucose uptake was reduced in *Col1A1*(Y/Y) MEFs compared to controls, and we also observed a reduction in pyruvate and lactate production into the medium, confirming defects in glucose utilization (Fig. 4d). Similarly to what was observed in the NMR analysis of the liver (Suppl. Fig. 3c), CCNY overexpression also led to decreased intracellular levels and deficient uptake of several amino acids (Suppl. Fig. 4a,b).

**Fig. 4.**
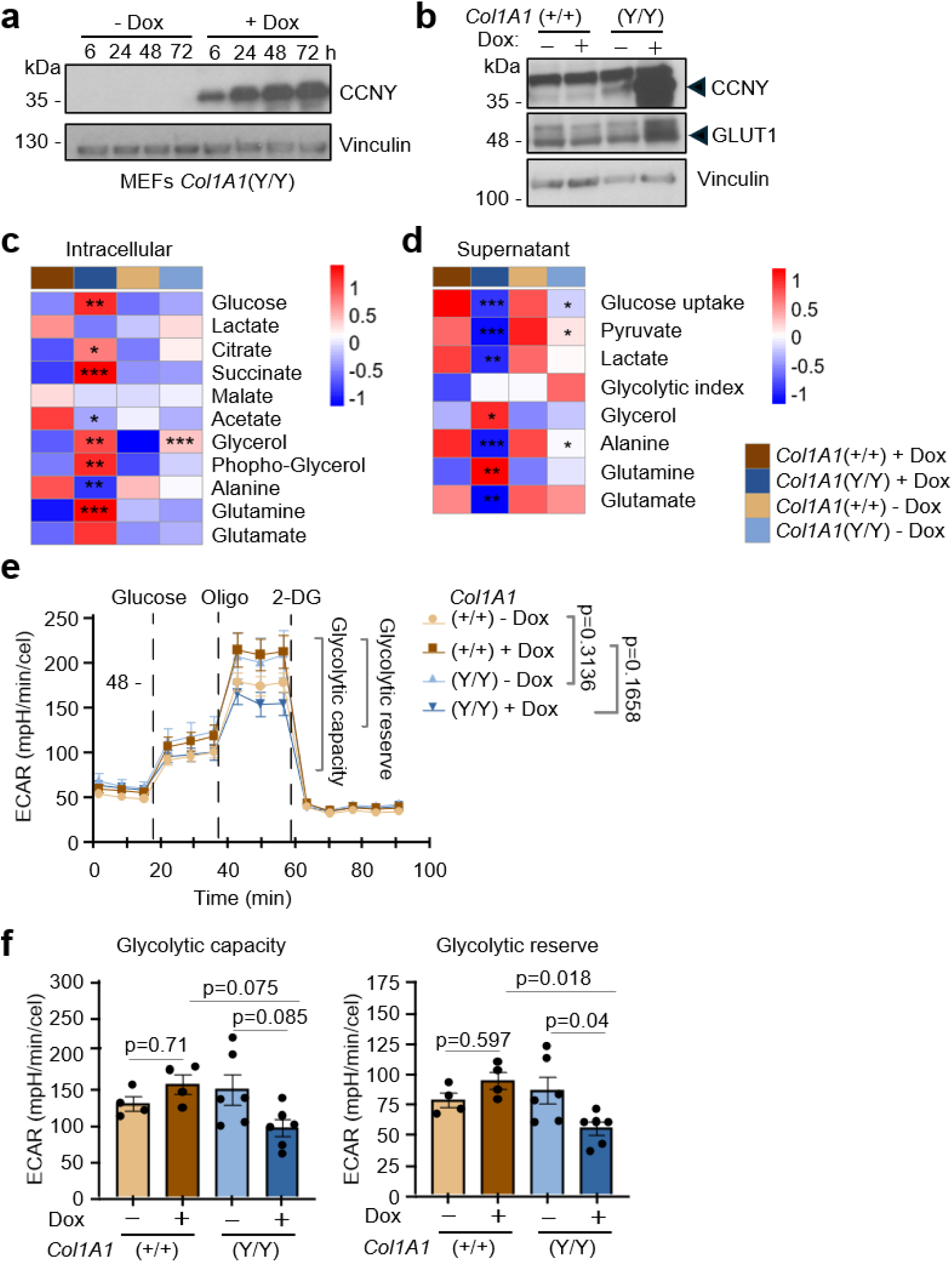
Glucose metabolism alterations in CCNY-overexpressing fibroblasts. **a)** Immunoblot of CCNY in mouse embryonic fibroblasts (MEFs) from *Col1A1*(Y/Y) mice treated with 1 μg/mL Dox at the indicated time points. **b)** Immunoblot of CCNY and GLUT1 in *Col1A1*(+/+) and*Col1A1*(Y/Y) MEFs untreated or treated with Dox. Black arrows indicate the specific bands corresponding to each protein. In a,b, vinculin was used as a loading control. **c,d)** Heatmap of z-score normalized data from NMR metabolite concentrations measured in intracellular polar extracts (**c**) and cell media (**d**) of MEFs using ^13^C-Glucose and treated with or without Dox. n=4-6 cultures per sample. Data were analyzed using two-tailed unpaired Student’s t-test of *Col1A1*(Y/Y) vs. control MEFs with or without Dox (*p<0.05; **p<0.005; ***p<0.0005). **e)** Seahorse analysis of *Col1A1*(+/+) and *Col1A1*(Y/Y) MEFs treated with or without Dox. Time course analysis of extracellular acidification rate (ECAR) with the specific inhibitors administrated in each experiment. Data is expressed as means ± SEM and statistics were analyzed with two-way ANOVA with Šidák’s multiple comparison test of compared conditions. (n = 4-6 cultures per group). **f)** Glycolytic capacity and the glycolytic reserve in the samples indicated in the previous assay. Data are represented as means ± SEM and were analyzed by one-way ANOVA with Tukey’s multiple comparisons.

We also performed Seahorse analysis in MEFs to test the oxidative phosphorylation (OXPHOS) capacity of their mitochondria and the glycolytic rate. We did not get differences in OXPHOS parameters (basal respiration, ATP-linked respiration, proton leak, maximum respiration and spare capacity) upon CCNY induction in MEFs (Suppl. Fig. 4c). The extracellular acidification rate (ECAR), however, was altered in CCNY-overexpressing cells (Fig. 4e), with a tendency to reduced glycolytic capacity and significant defects in the glycolytic reserve in these mutant cells (Fig. 4f), suggesting that CCNY-overexpressing cells are already at maximum capacity showing a cell-autonomous limitation in their metabolic responses.

### Proteomic analysis of CCNY-overexpressing livers suggests new interacting partners and regulatory alterations in glucose metabolism

To obtain further insights into possible mechanisms of action behind these observations, we performed proteomic studies with FLAG immunoprecipitates. *Col1A1*(Y/Y) and control *Col1A1*(+/+) mice were treated during 6 days with Dox diet, and liver lysates were used for FLAG pulldown and mass spectrometry analysis (Fig. 5a). Using a threshold of fold change (FC) higher than 3 and a p-value < 0.05 we obtained 678 proteins over-represented in FLAG-IP lysates from CCNY overexpressing mice, being CCNY the most over-represented protein (Fig. 5b). Several CDKs were found among the over-represented proteins in the FLAG IPs including CDK1, CDK2, CDK4, CDK6, and CDK18; Fig. 5b). Apart from a correlation with CDK4 in liver cancer (Chen *et al*, 2020), and the known binding of Y-type cyclins to members of the CDK14-CDK18 subfamily (Mikolcevic *et al*., 2012a), these cell cycle CDKs have never been described as partners of cyclin Y. We could confirm the interaction between cyclin Y-FLAG and CDK1 or CDK6 using FLAG IP in liver lysates from *Col1A1*(+/+), *Col1A1*(+/Y), and *Col1A1*(Y/Y) mice after 6 days of doxycycline diet (Fig. 5c), whereas other interactions were unclear due to poor efficiency of antibodies. We also confirmed the interaction between CCNY and CDK1, CDK2 and CDK17 in similar biochemical assays from CCNY-overexpressing MEFs, whereas CDK6 and CDK18 were not clearly detected in these cells (Suppl. Fig. 5a).

**Fig. 5.**
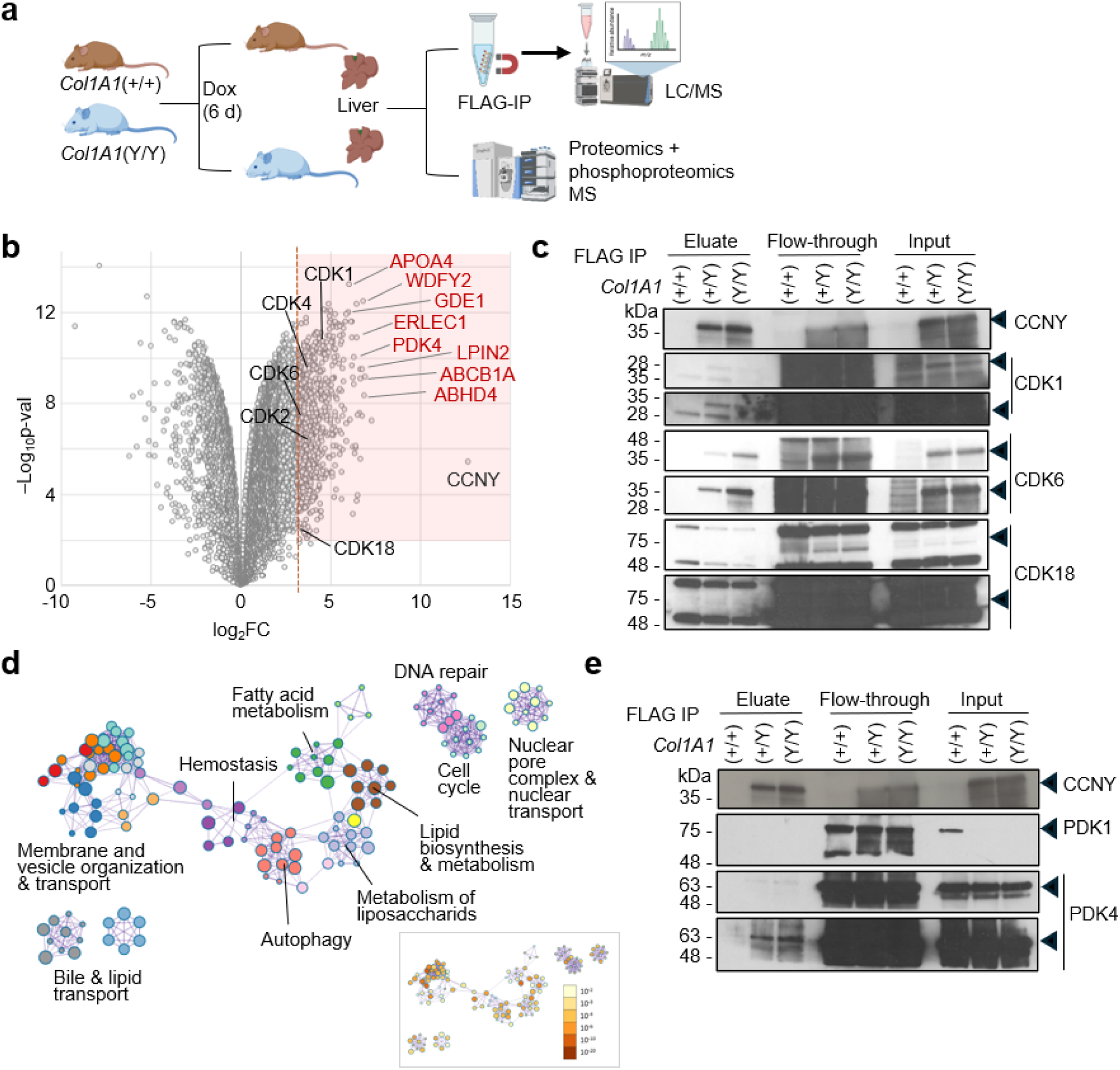
Proteomic and biochemical analysis of CCNY-FLAG complexes. **a)** Schematic representation of the immunoprecipitation (IP) couples to mass spectrometry (LC/MS), and proteomics/phosphoproteomics experimental workflow. **b)** Volcano plot of proteins over-represented in FLAG IPs in the IP-LC/MS experiment, with proteins showing fold change (FC) > 3 and p-value < 0.05 considered significant (red shade). Overrepresented CDKs are annotated in black font. Over-represented glucose metabolism proteins are annotated in red. **c)** FLAG pull-down of liver samples from *Col1A1*(+/+), *Col1A1*(+/Y) and *Col1A1*(Y/Y) mice treated 6 days with Dox diet. The noted CDKs were immunoblotted in the eluate, flow-through and input fractions. **d)** Metascape analysis of the network of pathways enriched in proteins over-represented in panel b). **e)** Immunodetection of the indicated proteins in similar FLAG-pulldowns as in c). In c,e, black arrows indicate the specific bands corresponding to each protein.

Several key metabolic regulators were found among the most enriched proteins in CCNY-FLAG immunolysates, including proteins involved in lipid transport such as Apolipoprotein A4 (APOA4) or ABCB1A, or lipid metabolism, including ABHD4, GDE1 or LPIN2 (Fig. 5b,d). Two of these top over-represented proteins were involved in glucose metabolism, including WDFY2, a poorly-characterized protein involved not only in the endosomal control of hepatic AKT2 signaling, but also in insulin-stimulated AKT2 phosphorylation and glucose uptake (Zhang *et al*, 2020b), and PDK4, a key regulator of glucose and fatty acid metabolism and homeostasis via phosphorylation of the pyruvate dehydrogenase subunits (Fig. 5b). Additional proteins involved in glucose metabolism such as PDK3, MTORC1, OGDH, and PFKL, the rate limiting step in glycolysis, were also enriched in CCNY-FLAG immunoprecipitates (Suppl. Table 1). We confirmed the interaction between CCNY-FLAG and PDK4 (Fig. 5e), whereas we observed no specific interaction with PDK1. In addition, PDK1 levels were downregulated in *Col1A1*(+/Y), and *Col1A1*(Y/Y) livers both at the protein (Fig. 5e) and mRNA levels (Suppl. Fig. 5b). Altogether, these data suggested the interaction between CCNY and a large networks of proteins involved in membrane and vesicle organization and transport, lipid metabolism, DNA repair and cell cycle, among others (Fig. 5d, Suppl. Fig. 5c,d and Suppl. Table 1).

We also submitted liver samples from control and *Col1A1*(Y/Y) overexpressing mice to proteomics + phospho-proteomics analysis (Fig. 5a). PCA and clustering analysis of total protein levels separated the two genotypes with further separation by sex (Suppl. Fig. 6a,b). CCNY was the most upregulated protein, and additional cyclins and CDKs upregulated included CCND1, CDK1, CDK17 and CDK11B (Fig. 6a, Suppl. Fig. 6c and Suppl. Table 2).

**Fig. 6.**
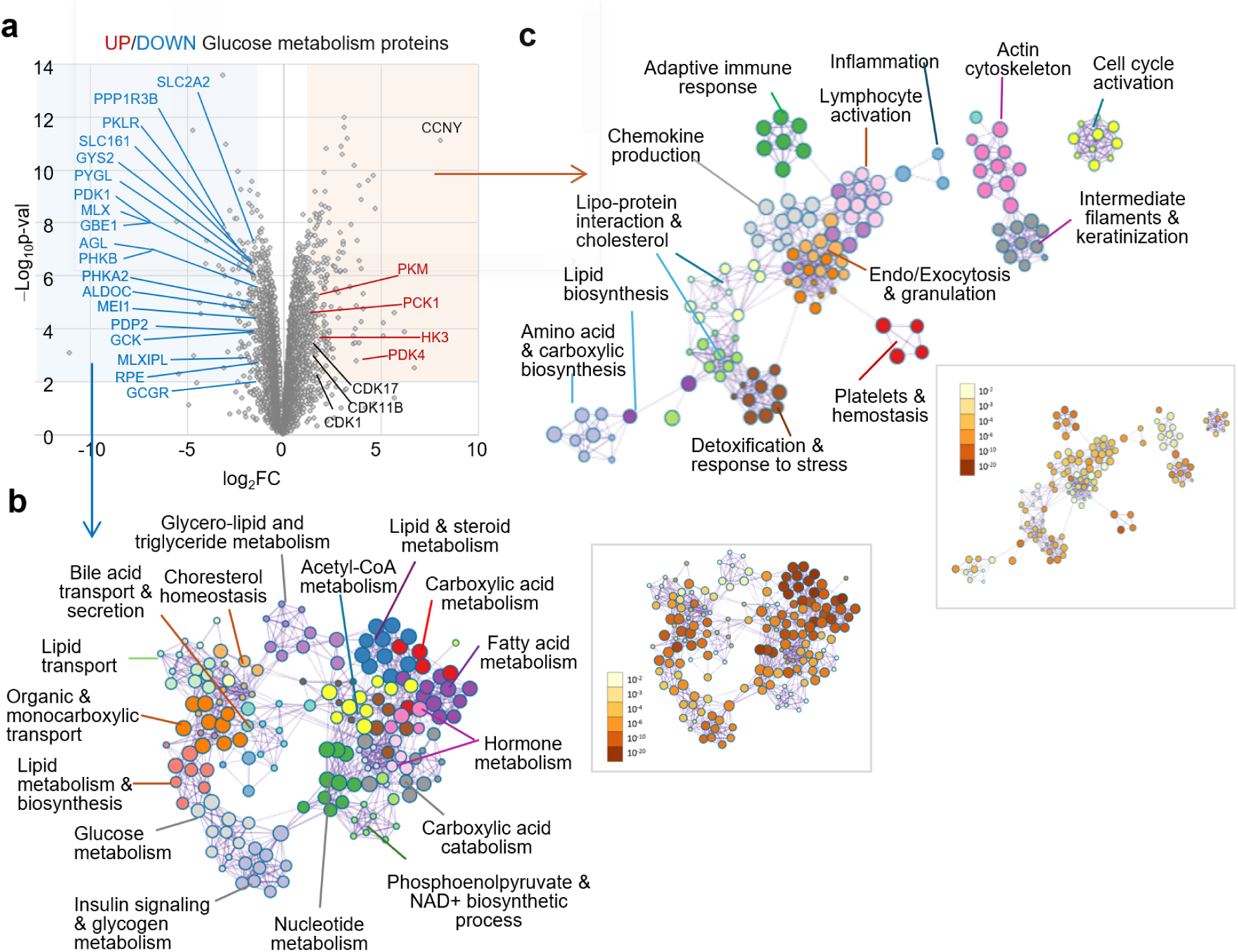
Total protein levels in CCNY-overexpressing livers. **a)** Volcano plot of proteins upregulated in CCNY-overexpressing livers. Proteins with logFC>1 or <−1 and p-value <0.05 were select as upregulated (red) or downregulated (blue), respectively. Upregulated CDKs are shown in black font. Up- (red) or down-(blue) regulated proteins involved in glucose metabolism are also indicated. **b)** Main pathways representing proteins downregulated at the total protein level in *Col1A1*(Y/Y) livers. Selected names group individual pathways (dots) with proteins in common. The p-value of these pathways is shown as in the panel to the right (Metascape analysis). **c)** Main pathways representing proteins upregulated at the total protein level in *Col1A1*(Y/Y) livers. Selected names group together individual pathways (dots) with proteins in common. The statistical analysis of these pathways is color-coded in the panel to the right (Metascape analysis).

Most of the downregulated proteins in the liver from CCNY-overexpressing mice were involved in the metabolism of carboxylic acids and glucose, as well as transport and secretion of molecules (Fig. 6a,b), including GLUT2 (also known as SLC2A2, the main glucose transporter of hepatocytes), GCGR (glucagon receptor), and PDK1, thus confirming previous observations in immunoblots and mRNA levels (Fig. 5e and Fig. 6a). Four proteins involved in glucose metabolism were upregulated: HK3 (Hexokinase 3), PKM (Pyruvate Kinase M), PCK1 (Phosphoenolpyruvate carboxykinase 1) and PDK4 (Fig. 6a), which we validated as a CCNY partner (Fig. 5e). Pathway analysis of upregulated proteins revealed the upregulation of pathways related to lipid and cholesterol synthesis, stress response, alterations in cytoskeleton and cell cycle activation (Fig. 6c). Interestingly, we detected overexpression of similar pathways (e.g. lipid and carboxylic acid biosynthesis, hormone biosynthesis and bile acid secretion and circulation) in liver hepatocellular carcinomas with CCNY overexpression compared to similar tumors with low levels of this cyclin (Suppl. Fig. 6d).

We next focused on the alteration of phospho-sites in 3 male and 3 female CCNY-overexpressing livers compared to littermate controls (Fig. 5a). After quality filters (see Methods) we quantified 17,694 phosphosites in 3,702 unique proteins. Principal component analysis (Fig. 7a) and clustering (Fig. 7b) of phospho-proteomics data indicated similar profile among all control samples whereas *Col1A1*(Y/Y) male and female samples diverged apart. These sex-based differences in CCNY-overexpressing mice were mostly related to over-representation of lipid-related pathways among the changes observed in females versus males (Suppl. Fig. 7a). After correcting phosphosite abundance with total protein level, 386 phosphosites were upregulated (logFC>=2) and 291 downregulated (logFC<=2; Fig. 7c and Suppl. Table 3). Among them, CCNY Ser100 phosphorylation, a known target of CDK16 (Shehata *et al*, 2015), was upregulated; whereas phosphorylation of CCNY in Ser326, which is mediated by AMPK activity and promotes autophagy (Dohmen *et al*, 2020), was among the top downregulated hits (Suppl. Fig. 7b,c).

**Fig. 7.**
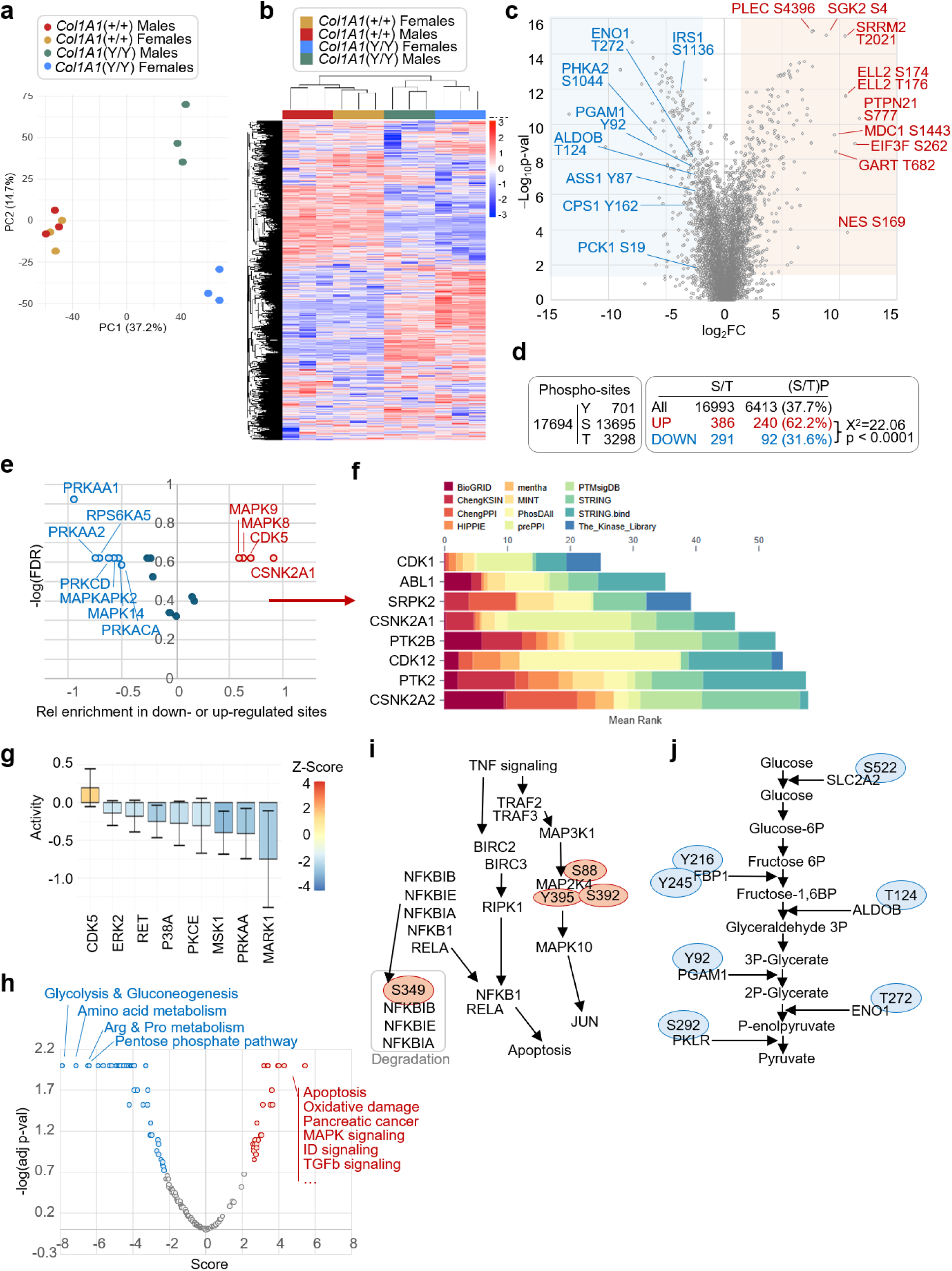
Phospho-proteomics analysis of CCNY-overexpressing livers. **a)** Principal component analysis (PCA) of z-score corrected phosphosite value from three mice per genotype and gender. PC1 and PC2 variances are represented in percentage. **b)** Clustered heatmap of z-score corrected data from phosphosite values (rows) in the indicated groups (columns). **c)** Volcano plot of phosphosites upregulated in CCNY-overexpressing livers. Sites with logFC >2 or <−2 and p-value <0.05 were selected as upregulated (red) or downregulated (blue), respectively. The top upregulated (red) and downregulated (blue) phosphosites in proteins involved in glucose metabolism (blue) are indicated. **d**) Quantification of phosphosites (Y, S and T) detected in the analysis of *Col1A1*(Y/Y) and control livers and number and percentage of S or T sites followed by a P, among the total S/T sites or those up- or down-regulated in CCNY-overexpressing livers. Differences were analyzed using Chi-square. **e**) Kinase Substrate Enrichment Analysis (KSEA) showing the relative enrichment of kinase substrates among the upregulated (red) or downregulated (blue) phosphosites. **f**) Kinase Enrichment Analysis (KEA3) to infer upstream kinases whose putative substrates are overrepresented proteins with upregulated phospho-sites. The plot shows the mean rank score in the indicated libraries with CDK1 as the top kinase. **g**) Robust Inference of Kinase Activity (RoKAI) analysis showing CDK5 as the only kinase significantly enriched as kinase responsible for the upregulated phosphosites in CCNY-overexpressing livers. **h**) Posttranslational Modification Substrate Enrichment Analysis (PTM-SEA) of main pathways representing upregulated (red) or downregulated (blue) phosphopeptides in CCNY-overexpressing livers. The top pathways are indicated and the complete list is provided as Suppl. Table 5. **i,j**) Schematic representation of upregulated (red) or downregulated (blue) phosphopeptides in the first overactivated (Apoptosis, **i**) or inhibited (Glycolysis and gluconeogenesis, **j**) pathways from the PTM-SEA analysis.

An initial analysis of the primary sequence of the phosphosites revealed that upregulated phosphosites were significantly enriched in (S/T)P sequences (62.2% of upregulated phosphopeptides compared to 31.6% of downregulated phosphopeptides; Fig. 7d). Although we cannot discriminate between direct targets and proteins dysregulated as a consequence of the primary targets, these findings are in agreement with over-activation of CDKs, which preferentially phosphorylate Proline-directed S/T sites. Within the top 10 sites hyperphosphorylated upon cyclin Y induction, 5 were S/T proline-directed sites, which is the consensus motif of CDKs: PTPN21, EIF3F, MDC1, Nestin (NES) and SGK2 (Fig. 7c). PTPN21, EIF3F and SGK2 have been previously described to regulate lipid and glucose metabolism (Gotoh & Negishi, 2015; Lian *et al*, 2024; Zhou *et al*, 2025). Subsequent studies using Kinase Substrate Enrichment Analysis (KSEA; Fig. 7e), Kinase Enrichment Analysis (KEA3; Fig. 7f) and Robust Inference of Kinase Activity (RoKAI; Fig. 7g) suggested CDKs, and in particular CDK1 and CDK5 (the CDK family member closest to atypical CDK14-18 CDKs) as main kinases responsible for upregulated phosphorylations in *Col1A1*(Y/Y) overexpressing livers. Posttranslational Modification Substrate Enrichment Analysis (PTM-SEA) also confirmed CDK5 as main driver of deregulated phosphoresidues (Supp. Table 4). On the other hand, AMPK (PRKAA) and p38α (MAPK14) kinases, as well as its main downstream target MK2 (MAPKAPK2), were consistently found enriched as kinases responsible for downregulated phospho-sites (Fig. 7e and Fig. 7g).

PTM-SEA analysis of the main cellular pathways dysregulated in CCNY-overexpressing livers at the level of phosphorylation suggested multiple phosphosites increasing apoptosis and oxidative damage in CCNY-overexpressing cells (Fig. 7h,i and Suppl. Table 5). Downregulated phosphorylations on the other hand mainly affected metabolic pathways, including glucose and amino acid metabolism, and the pentose phosphate pathway as top hits (Fig. 7h,j and Suppl. Table 5). In particular, some of the top phospho-residues downregulated after normalizing by protein content controlled critical steps in glycolysis, including ALDOB T124 or ENO1 T272 (Fig. 7c and Fig. 7j, in addition to T522 phosphorylation in SLC2A2 (Solute Carrier Family 2 Member 2; Fig. 7j), also known as Glucose Transporter Type 2 (GLUT2), a key protein responsible for moving glucose across cell membranes, particularly in the liver, in agreement with the metabolic defects observed in *Col1A1*(Y/Y) mice.

## Discussion

CCNY and its homolog Cyclin Y-like 1 (CCNYL1) are atypical cyclins defined by a single cyclin box and unique N-terminal lipidation motifs. These Y-type cyclins bind and activate certain CDKs, notably CDK14 and CDK16, often targeting complexes to the plasma membrane (Martinez-Alonso & Malumbres, 2020; Mikolcevic *et al*., 2012a; Quandt *et al*., 2020). While initially recognized for roles in cell cycle control and WNT signaling, emerging evidence indicates CCNY/CCNYL1 can also influence metabolic processes. Specifically, *Ccny* ablation in the mouse leads to decreased body weight and adiposity (An *et al*., 2015). These mutant animals resist high-fat diet–induced obesity and have impaired white adipose tissue (WAT) development, suggesting CCNY acts downstream of pro-adipogenic signals and its loss biases metabolism toward energy expenditure over storage. *Ccny*-deficient mice also display higher blood glucose levels and hepatic insulin resistance (An *et al*., 2015), although the function of cyclin Y in these pathways remains unclear.

In vivo genetic studies show that Cyclin Y is essential for WNT signaling in early development; for example, *Xenopus* and zebrafish embryos require maternal Cyclin Y for LRP6 phosphorylation and WNT target gene expression (Niehrs & Acebron, 2012). In mammals, *Ccny/Ccnyl1* double-knockout is embryonic lethal by E16.5 due to failure of WNT-driven proliferation in multiple stem cell compartments (Zeng *et al*., 2016). Interestingly, perturbations in WNT signaling are known to affect metabolism. LRP6 mutations in humans cause metabolic syndrome characterized by hyperlipidemia and glucose intolerance (Jeong & Jho, 2021). Overall, CCNY’s position upstream of WNT/β-catenin and GSK-3β could link this cyclin to pathways that regulate not only cell fate but also systemic metabolism. WNT-mediated inhibition of GSK-3β a key inhibitor of glycogen synthesis, could promote for instance glycogen and lipid synthesis (Xue *et al*, 2025).

Strikingly, in this work we found that sustained CCNY overexpression from the Rosa26(rtTA); *Col1A1*(Y) alleles in the presence of Dox proved toxic, causing premature death in adult mice (Fig. 1). The phenotype suggests pathological hypoglycemia accompanied by a starvation response (low insulin and elevated glucagon) accompanied by liver damage. The fact that CCNY-overexpressing mice also showed lower glucose rise in pyruvate tolerance tests suggests a defect in hepatic gluconeogenesis, in agreement with the abnormal accumulation of elevated levels of major circulating gluconeogenic substrates, such as lactate, alanine or glutamine (Fig. 3). This phenotype mimics the Warburg Effect seen in aggressive cancers, characterized by systemic glucose consumption and high lactate production, to increase demand of gluconeogenesis intermediates (Chen *et al*, 2024; Holeček, 2023; Luo *et al*, 2023).

Proteomic and phospho-proteomic analysis of CCNY-overexpressing cells suggested physical and functional interaction between CCNY and several critical regulators of metabolism. CCNY immunoprecipitates were enriched in pyruvate dehydrogenase kinase isoforms PDK3 and PDK4, mTORC1 (mTOR complex 1), OGDH (α-oxoglutarate dehydrogenase, a TCA cycle enzyme that converts α-ketoglutarate to succinyl-CoA and controls TCA cycle throughput), and PFKL (phosphofructokinase liver type, a key glycolytic enzyme rate-limiting for glycolysis). All five factors are central to glucose metabolism. When considering total protein levels and changes on phosphorylation (after normalization for protein levels), the loss of metabolic control is the most striking molecular finding among downregulated pathways. In particular, the downregulation of phosphorylation on glycolytic and gluconeogenic enzymes, amino acid biosynthesis and related metabolic routes confirms the liver has lost its capacity to respond to nutrient cues creating a “metabolic zombie” state. Apoptosis as the top upregulated pathway at the phospho-proteomic level suggesting that the starving signals in the liver are ultimately triggering pro-death signaling. Altogether, whereas CCNY deficiency skews toward energy expenditure (lean mice with active BAT) (An *et al*., 2015), CCNY excess causes glucose deregulation and energy deficit (weight loss, hypoglycemia) despite attempts to increase intake. We hypothesize that CCNY normally promotes energy storage and glucose utilization efficiency and is part of a hepatic pacemaker on gluconeogenesis. When CCNY is absent, animals are inefficient in storing energy (leading to resistance to obesity), and when CCNY is too high, energy usage becomes uncoupled from proper nutrient balance (leading to wastage and hypoglycemia).

What is the critical CDK-partner for this function is currently unclear. Our data suggests a statistically significant enrichment of Serine/Threonine-Proline (S/T-P) sites, the classic consensus for CDKs and MAPKs, among the upregulated phosphosites. Phospho-site analysis in CCNY-overexpressing livers suggest CDK5 (the closest CDK to the CDK14-18 subfamily) or CDK1 as responsible kinases (Fig. 7), although it is difficult to assign this to specific family members due to the promiscuity in their consensus phosphorylation sites, and the poor representation of other CDKs in these databases. The fact that CCNY is overexpressed in our model may generate non physiological complexes given the promiscuity within the CDK and cyclin families (Malumbres, 2014). Multiple CDKs, including at least CDK1, CDK2, CDK4, CDK6 and others, have been shown to impact cell metabolism at different levels (Salazar-Roa & Malumbres, 2017). Cyclin D1-CDK4 and CDK1 have been described to inhibit gluconeogenesis in hepatocytes (Gassaway *et al*, 2019; Lee *et al*, 2014), and CDK4 to potentiate OXPHOS and glucose uptake in muscle (Bahn *et al*, 2023). Cyclin D3-CDK6 complexes on the other hand phosphorylate intermediates of glycolysis redirecting those to pentose phosphate pathway and serine pathways in cancer cells to produce antioxidant species counteracting reactive oxygen species production (Wang *et al*, 2017). Our data suggest that overexpressed CCNY may bind several of them, in addition to known partners of the CDK14-CDK16 subfamily, at least when expressed at high levels. It is tempting to speculate that CCNY may link metabolism and cell cycle regulation in specific situations, as previously proposed for the connection between WNT signaling and cell cycle progression specifically by CCNY-CDK14 complexes (Davidson & Niehrs, 2010; Niehrs & Acebron, 2012).

## METHODS

### Generation of mouse models

For the generation of the conditional knock-in mouse model, the human *CCNY* cDNA was fused to 3X FLAG sequence in the C-terminal and cloned into pBS31 vector (Beard *et al*., 2006) by Mlu1 restriction enzyme. Cloned vector (20 μg) was electroporated in KH2 ES cells carrying the M2-rtTA gene inserted within the Rosa26 allele, together with a plasmid expressing the Flp recombinase (pCAG-Flp, 50 μg) using the Bio-Rad Gene Pulser II. Flp-mediated recombination at the modified *Col1A1* locus of the KH2 cells resulted in the insertion of a cassette containing the *hCCNY*-FLAG cDNA under the control of the Dox-responsive promoter (TRE) was inserted downstream of the *Col1A1* gene polyA, and positive clones were selected in 200 μg/mL of Hygromycin. Germ line chimeras were obtained from positive clones by microinjection in CD1 8-cell embryos and bred to C57BL6 females. Mice were housed in the pathogen-free animal facility of the CNIO (Madrid) and maintained under a standard 12-h light-dark cycle, at 23°C with free access to food and water. Mice were fed with special diet containing 2g doxycycline hyclate/kg of diet when indicated. Routine genotyping of the *CCNY* alleles was done by PCR using DNA from the ear of the mice and the oligonucleotides indicated in Suppl. Table 6. All animal work and procedures were approved by the ISCIII committee for animal care and research and were performed in accordance with the CNIO Animal Care program.

### Metabolic and behavior assays

For metabolic studies, mice were housed individually in experimental metabolic cages three days prior to the start of measurements, allowing for parameter adjustment and acclimation to the new environment. The experiment was performed in an OXYLETPRO-PHYSIOCAGE (Panlab). Intake and activity were measured each 20 minutes, meanwhile oximetry was assessed each 10 minutes. Food and water intake was determined by high precision extensometer weight transducers. Activity and rearing were measured through platform sensors which transduced weight changes into activity, and infrared sensor to detect the occurrence and duration of rearing. Finally, O_2_ and CO_2_ volumes were measured using LE405 sensors, determining the volume of O_2_ consumed, volume of CO_2_ produced and the energy exchange. Mice were monitored for 72 hours, and subsequent data analysis was performed using RStudio.

For the Open Field experiment, 7-week-old male and female mice were evaluated 4 days after the start of doxycycline diet. Mice were placed in the behavior testing room one day prior the start of the experiment, to allow acclimatation. Males were assessed first to prevent any potential behavioral interference caused by residual female odor. Testing was conducted in a transparent polyvinyl chloride (PVC) box (A 50 x 50 x 41 cm). Each mouse was recorded for 15 minutes, and the arena was cleaned and dried between animals. Analysis of the experiment was carried out using MatLAB software and following the pipeline described in (Zhang *et al*, 2020a).

For Rotarod assays, 8-weeks-old male and female mice were evaluated in a Rotarod LE8200 (Panlab). For two days, mice underwent training, which involved three consecutive attempts per day, separated 5 minutes between them. In each trial, mice were placed over the running wheel at low speed (4-8 revolutions per minute (rpm) for a maximum of 1 minute). The test phase consisted in an accelerating protocol, with the speed increasing from 4 to 40 rpm over 5 minutes. Mice were evaluated twice per day during 4 consecutive days; the mean values for each mouse were plotted. Mice were treated with doxycycline diet since the beginning of the training.

### Quantification of blood glucose and metabolites and other blood parameters

Blood glucose level was determined using glucose strips and glucometer (Accu-Chek, Roche, #07819382037). For glucose tolerance test (GTT) and pyruvate tolerance test (PTT), mice under doxycycline diet for 6 days were subjected to 16 hour-fasting (6 pm-10 am). Following fasting, mice were intraperitoneally injected either with 1 g/Kg D-Glucose (Sigma #G7528) for GTT or with 1 g/Kg sodium pyruvate (Merck, #P256) for PTT, diluted in saline solution.

For other metabolites and blood parameters, blood was collected from heart of anesthetized mice in EDTA-coated tubes. Alanine aminotransferase, bile acids, albumin, blood urea nitrogen, alkaline phosphatase, γ-glutamyl transferase and cholesterol were measured using VetScan mammalian liver profile rotors (Abaxis #500-0040-12). Plasma was harvested by centrifugation of EDTA-coated blood collection tubes at 3.000 x g for 15 minutes at 4°C. Triglycerides (TG; #A11A01640) were measured using PENTRA 400 clinical chemistry analyzer (Horiba). Enzyme-linked immunosorbent assay (ELISA) was used to assess the plasma concentration of glucagon (Crystal Chem, # 81518) and insulin (Merck, #EZRMI). Assays were conducted following manufacturers’ instructions. Measurements were performed using 10 μl of plasma per well, and absorbance was measured at 450 nm using a microplate reader (CLARIOstar).

### Cell culture

Mouse embryonic fibroblasts (MEF) were obtained from extraction and physical disaggregation of E13.5 embryos. The liver and the head were removed to avoid contamination of differentiated hepatocytes and hematopoietic cells. MEFs were genotyped to confirm the presence of the desired alleles. Cell lines and primary cultures were tested for mycoplasma periodically. MEFs were cultured with DMEM-High glucose, 1% of GlutaMAX (Thermo Fisher Scientific, #35050061) and 10% of FBS and maintained at 37°C with 5% CO_2_ saturation.

### NMR measurements

For analysis of cell media metabolites, 500 μl of cell media supernatant was sampled for every time point and centrifuged at 10,000 x g for 10 min using 10 kD Spin Columns (Abcam, #ab93349) to eliminate debris and high molecular mass components. NMR samples were prepared mixing 400 μl of medium filtrate with 200 μl of NMR buffer (PBS 3X in 30% deuterated water, 0.3 mM TMSP, and 0.1% sodium azide) and transferred to 5 mm NMR tubes for measurement. For cell and tissue extracts NMR samples were produced in the same way after resuspending the polar and apolar dried fractions in polar buffer (PBS in 100% deuterated water, 0.3 mM TMSP, and 0.03 % sodium azide), or organic biphasic buffer (500 μl of CDCl_3_ plus 300 μl of 20 mM deuterated EDTA pH 6.0 in D_2_O: deuterated methanol (1:2)) and transferred to 5 mm NMR tubes for measurement. Liver extracts were resuspended in 600 μl of polar buffer and measured in 5 mm tubes. Cells and snap-frozen tissue samples were subjected to apolar/polar phase extraction. Briefly, cells plates were washed thoroughly with saline and excess liquid removed before adding cold methanol craping cells off and transferring them to the glass tubes. For tissue, 200 mg of liver/muscle was placed in a Precellys homogenization tube with 3 zirconia beads and 1 ml of cold methanol and 3 rounds of homogenization (2 x 20 s x 5500 rpm) were applied. Once tissue/cells are homogenized, tissue samples are resuspended in a total volume of 3 ml of cold methanol and cells in 4 ml. 1 volume of HCCl_3_ is added to each sample and vortexed vigorously. 1 volume of ddH_2_O is added to the mixture, vortexed vigorously and centrifuged at 4000 x g and 4 °C for 20 minutes. Then, the polar and apolar fractions were carefully separated and subsequently dried by lyophilization (polar fraction) or vacuum centrifugation (apolar) using a GeneVac. This allowed simultaneous extraction of cellular lipids and water-soluble metabolites from cells or tissues, as described before (Bakiri *et al*, 2017; Petruzzelli *et al*, 2014).

The plasma was collected from blood of anesthetized mice by heart puncture. Blood was collected in EDTA-coated tubes and centrifuged for 15 minutes at 3000 x g and 4°C in a tabletop centrifuge. 100 μl of plasma was mixed with 100 μl ice-cold 2X PBS buffer pH 5.05 containing 20 mM KF and 0.06% NaN_3_ in deuterium oxide (D_2_O), centrifuged for 10 min at 16000 x g and transferred to a 3 mm NMR sample tube. Metabolite levels in plasma samples were determined from the integrals of the most resolved and largest signals of each metabolite (for example methyl groups of lactate, alanine and pyruvate, and H4 proton of glucose) in 1D ^1^H NMR spectra acquired with a transversal relaxation filter (CPMG of 200 ms, t=0.4 ms) that attenuates the fast relaxing signals of macromolecules (proteins, lipids and lipoproteins) signals and optimizes the NMR signals of low mass metabolites for their quantification (Beckonert et al., 2007). Thus, the obtained values of integrals are not absolute concentrations but relative concentrations in approximate μM concentrations, referred to an NMR calculated average glycemia of 5 mM for the control group.

High-resolution NMR spectra were registered on a Bruker Avance spectrometer operating at 16.4 T (proton Larmor frequency of 700 MHz) at 293 K using a RT TXI probe with pulsed field gradient capabilities and equipped with a BACS120 sample changer. 1D proton NMR spectra (1dH) of cell growth media, and cell and tissue extract were recorded using a NOESY pulse sequence (noesygppr1d in Bruker nomenclature) employing pulse field gradients during the mixing time of 10 ms and low power (50 db) presaturation of the residual water signal during the 2 s recovery delay between consecutive scans. Analogous parameters were used to record 13C-filtered 1D proton NMR spectra (1dHC), which allows selective observation of 13C-bound protons (from 13C-glucose and metabolic transformation products thereof). 1dHC spectra were recorded using the first increment of a 2D sensitivity-enhanced 1H-13C heteronuclear single quantum coherence experiment (2dHC, hsqcetgpsisp2 in Bruker nomenclature) (Schleucher *et al*, 1994; Willker, 1993) without 13C decoupling during the 1.6 s acquisition time. All 1dH and 1dHC free induction decays (FIDs) were processed with exponential multiplication (0.5 Hz line-broadening) before Fourier transformation and baseline corrected afterwards using Topspin2.1 (Bruker). 1H chemical shifts were referenced to internal TMSP (0.00 ppm). 2D sensitivity-enhanced 1H-13C heteronuclear single quantum coherence spectra (2dHC, hsqcetgpsisp2 in Bruker nomenclature) (Schleucher *et al*., 1994; Willker, 1993) were recorded for each extract sample (at 13C natural abundance for tissue extracts) with 13C decoupling during the 60 ms acquisition time and 1.2 s recovery delay. These polar/apolar spectra were heavily folded in the 13C dimension, with a spectral width of 32/40ppm centered at 62/68 ppm, with 256-512 accumulations (for cell and tissue extracts, respectively) per each of the 64/70 complex points in the indirect 13C dimension, amounting to 6-12 h of measuring time each. All 1dH free induction decays (FIDs) were processed with exponential multiplication (0.5 Hz line-broadening) before Fourier transformation and followed by baseline correction using Topspin2.1 (Bruker). The 2dHC spectra were processed with NMRPipe (Delaglio *et al*, 1995) using squared cosine window functions in both dimensions, and were visualized and analysed using nmrViewJ (Johnson, 2004). ^1^H/^13^C chemical shifts were referenced to internal TMSP (0/0 ppm). Metabolites in 1dH spectra were quantified from their signal/s integral/s, using integration regions of variable size that were manually defined to include all metabolite signals using the AMIX3.8 software (Bruker).

Approximate concentrations of most metabolites not linked to glucose were obtained from signal integrals of 1D spectral regions, normalizing to total extracted material and number of protons.

For ^13^C natural abundance of tissues and for intracellular ^13^C-Glucose derived relative metabolite concentration determination (Beckonert *et al*, 2007), intensities of signals in 2D ^1^H-^13^C spectra of polar/apolar fractions of the cell extracts were used, without correction for proton number or transfer function (reference (Saborano *et al*, 2019)). All these mathematical operations were performed on the 1dH raw data exported from AMIX (1D spectra) or from NMRView (2D spectra) and imported into Excel. Significant metabolite level changes were identified using p-values obtained with a two-tailed Student’s t test in Excel, which were then examined and plotted.

### Seahorse analysis

MEFs were treated with or without 1 μg/ml doxycycline for 4 days. The day prior to the experiment, MEFs were harvested and seeded at 8000 cells/well in Seahorse XFe96/XF Pro plates (Agilent, #103794-100). Confluency and distribution were checked. On the day of the experiment, cells were washed once, and media was replaced with Seahorse XF base medium (Agilent, #103335-100) supplemented with 10mM glucose, 1mM pyruvate and 2mM glutamine. For glycolytic assay the supplemented medium only contained pyruvate and glutamine. To test the mitochondrial respiratory capacity, the Seahorse XF Cell Mito Stress test kit was performed (Agilent, #103015-100). The electron transport chain (ETC) modulators were added to the media following a specific order: 1.5 μM oligomycin, 2 μM carbonyl cyanide 4-trifluoromethoxy phenylhydrazone (FCCP) and 0.5 rotenone/antimycin A (Rot/AA). The oxygen consumption rate (OCR) was measured in pmol/minute. To assess the glycolytic status of MEFs, Seahorse XF Glycolytic rate assay was performed. Different modulators were added following a specific order: 10 mM glucose (Sigma #G7528), 2 μM oligomycin (Merck #75351) and 50mM 2-deoxy-D-glucose (2-DG, Merck #D6134). The extracellular acidification rate (ECAR) was measured in mpH/minute. After the experiment, plates were fixed with 4% PFA for 15 minutes at RT. Then, nuclei were stained for 30 minutes with 1 μg/ml DAPI, and cell number were quantified using Opera Phenix and Harmony software. Normalization of each well was done. The analysis of the experiment was done using the Seahorse Analytics software (Agilent). Wells with injection problems or abnormal response to modulators were removed from the final analysis.

### RT-qPCR

Total RNA was extracted from mouse liver using Trizol (Sigma, #T9424), according to the manufacturer’s instructions. Reverse transcription from 1 μg of total RNA was performed with the M-MLV Reverse Transcriptase, RNase H minus, enzyme (Promega, #M3682), followed by quantitative PCR (qPCR), using QuantStudio™ 6 Flex Real-Time PCR System (Applied Biosystems). Relative quantification of mRNA expression was performed by real-time PCR using primers against Pdk1 (Fw: CGGGCCAGGTGGACTTCTA; Rv: AGGCAACTC TTGTCGCAGAA) and Rn-18s-rs5 (Fw: GGCTCTTCCGTGTCTACGAG; Rv: CCAGCCAACGTAGAAAAGCC). Data were analyzed using the comparative delta–delta Ct method and are expressed as the ratio between the expression of Pdk1 and the housekeeping gene Rn-18s.

### Biochemical and proteomics studies

For immunoprecipitation experiment, proteins were eluted by incubating FLAG-beads (Merck, #M8823) for 15 minutes at 95°C 1400 RPM in 2% SDS, 100 mM TEAB pH 7. Proteins were reduced and alkylated (15 mM TCEP, 25 mM CAA) 1 h at 45 °C in the dark. Then, samples were digested following the solid-phase-enhanced sample-preparation (SP3) protocol (Hughes *et al*, 2019). Briefly, ethanol was added to the samples to a final concentration of 70% and proteins were incubated for 15 minutes with SP3 beads at a bead/protein ratio of 10:1 (wt/wt). Beads were rinsed using 80% EtOH and proteins were digested with 100 μl of trypsin (Promega, 0.25 μg per sample) in 50 mM TEAB pH 7.5 16 h at 37°C. LC-MS/MS was done by coupling an UltiMate 3000 *RSLCnano* LC system to a Orbitrap Exploris 480 mass spectrometer (Thermo Fisher Scientific). Peptides were loaded into a trap column (Acclaim™ PepMap™ 100 C18 LC Columns 5 μm, 20 mm length) for 3 minutes at a flow rate of 10 μl/min in 0.1% FA. Then, peptides were transferred to an EASY-Spray PepMap RSLC C18 column (Thermo) (2 μm, 75 μm x 50 cm) operated at 45°C and separated using a 60 min effective gradient (buffer A: 0.1% FA; buffer B: 100% ACN, 0.1% FA) at a flow rate of 250 nl/min. The gradient used was from 2% to 6% of buffer B in 2 min, from 6% to 33% B in 58 minutes, from 33% to 45% in 2 minutes, plus 10 additional minutes at 98% B. Peptides were sprayed at 1.5 kV into the mass spectrometer via the EASY-Spray source and the capillary temperature was set to 300 °C. The mass spectrometer was operated in a data-independent acquisition (DIA) mode using 60,000 precursor resolution and 15,000 fragment resolution. Ion peptides were fragmented using higher-energy collisional dissociation (HCD) with a normalized collision energy of 29. The normalized AGC target percentages were 300% for Full MS (maximum IT of 25 ms) and 1000% for DIA MS/MS (maximum IT of 22 msec). 8 m/z precursor isolation windows were used in a staggered-window pattern from 396.43 to 1004.70 m/z. A precursor spectrum was interspersed every 75 DIA spectra. The scan range of the precursor spectra was 390-1000 mz. Raw files were processed with DIA-NN (version 1.8.2) using the library-free setting against a mouse protein database (UniProtKB/Swiss-Prot, One Protein Sequence Per Gene, 21,990 sequences). Precursor m/z range was set from 390 to 1010, and all other settings were left at their default values. A protein group intensity table was obtained by summing the precursor quantity values from the table report.tsv after filtering for a Lib.Q.Value and Lib.PG.Q.Value < 0.01. The protein group intensity table was loaded in Prostar (v1.26.4) (Wieczorek *et al*, 2017) for further statistical analysis. Briefly, samples were normalized using median intensity, proteins with less than four valid values in at least one experimental condition were filtered out. Missing values were imputed using the algorithms SLSA (Bø *et al*, 2004) for partially observed values and DetQuantile for values missing on an entire condition. Differential analysis was done using the empirical Bayes statistics Limma. Proteins with a p-value < 0.05 and a log2 ratio >3 defined as interactors. The FDR was estimated to be below 0.5% by Benjamini-Hochberg.

For proteomics and phosphoproteomic analysis Liver tissue samples were lysed in a FastPrep-24 ®5G (preset LIVER program) with 2.8 mm ceramic beads using 2% SDS, 100 mM TEAB pH 7.5 supplemented with 1:1000 (v/v) of benzonase (Novagen) and 1:100 (v/v) of HaltTM phosphatase and protease inhibitor cocktail 100X (Thermo Fisher Scientific). Samples were centrifuged at 10000 x g for 10 minutes. Protein concentration was determined using Pierce™ BCA Protein Assay. Then, samples (200 μg) were digested using on-bead protein aggregation capture (PAC) with ReSyn Biosciences MagReSyn® Hydroxyl microparticles (ratio Protein/Beads 1:2) in an automated King Fisher instrument (Thermo). Proteins were digested for 16 h at 37 C with LysC/Trypsin mix at a protein:enzyme ratio of 1:50 in 50 mM TEAB pH 8.0 (Trypsin Sequence grade, Sigma; LysC, Wako). After digestion, samples were Speed-Vac dried.1 μg of sample was reserved for the total proteome analysis. Phosphopeptides were enriched using MagReSyn Zr-IMAC HP beads following the automated phosphopeptide enrichment on KingFisher protocol (2). After enrichment, samples were acidified to a final concentration of 0.5% FA. LC-MS/MS was done by coupling an Evosep One system to an Orbitrap Exploris 480 mass spectrometer (Thermo Fisher Scientific). Peptides were loaded into Evotips (Evosep) and separated using the standard “Whisper” gradient with a throughput of 20 samples per day (SPD) on an Aurora Elite TS column of 15 cm length, 75 μm internal diameter (i.d.), packed with 1.7 μm C18 beads (IonOpticks). The column temperature was maintained at 55°C using a column heater (IonOpticks). Peptides were sprayed at 1.5 kV into the mass spectrometer via the EASY-Spray source and the capillary temperature was set to 280 °C. The mass spectrometer was operated in a data-independent acquisition (DIA) mode using 120,000 precursor resolution and 30,000 fragment resolution. Ion peptides were fragmented using higher-energy collisional dissociation (HCD) with a normalized collision energy of 25.9. The normalized AGC target percentages were 300% for Full MS (maximum IT of 25 ms) and 1000% for DIA MS/MS (maximum IT of 25 msec). 25.9 m/z precursor isolation windows were used from 469.5 to 1143.9 m/z. A precursor spectrum was interspersed every 25 DIA spectra. The scan range of the precursor spectra was 350-1400 mz. Raw files were processed with Spectronaut. A directDIA analysis was performed using the BGS Phospho PTM Workflow (phosphoproteome analysis). For the global protein analysis, a BGS Factory Workflow was used. A mouse protein database (UniProtKB/Swiss-Prot, One Protein Sequence Per Gene, 21,990 sequences) was used for the search. Protein LFQ method was set as QUANT 2.0. Cross-Run normalization, Major Group Top N and Minor Group Top N were unchecked. All other settings were left at their default values. A table containing the quantification of all the identified phosphorylation sites was loaded in Prostar (Wieczorek *et al*., 2017) for the phosphosite analysis. Briefly, phosphosites with less than four valid values in at least one experimental condition were filtered out. Missing values were imputed using the algorithms SLSA (Bø *et al*., 2004) for partially observed values and DetQuantile for values missing on an entire condition. Differential analysis was done using the empirical Bayes statistics Limma. Sites with a p-value < 0.05 and a log2 ratio >2 or <-2 were defined as regulated. The FDR was estimated to be below 5.06% by Benjamini-Hochberg. For the total protein analysis, the protein intensity table was loaded in Prostar and the same filtering and imputation settings were used. Differential analysis was done using the empirical Bayes statistics Limma. Proteins with a p.value < 0.05 and a log2 ratio >1 or <-1 were defined as regulated. The FDR was estimated to be below 5.22% by Benjamini-Hochberg.

### Protein immunodetection

For immunoblot analysis, cells were extracted on ice in RIPA lysis buffer (50 mM Tris-HCl pH 7.5, 10 mM EDTA, proteases inhibitor (Roche #05892970001), 1 mM sodium orthovanadate, 10 mM sodium fluoride, 10 mM β-glycerophosphate, 1mM dithiothreitol, 0.1% SDS, 0.5% NP-40, 150 mM sodium chloride). Cell lysates were disaggregated by mechanical forces (pipetting up-down repeatedly) and centrifuged at 13.200 rpm for 15 min at 4°C in a tabletop centrifuge. Protein concentration was quantified using the Bradford method (Bio-rad, #5000006). Protein extracts were mixed with RIPA buffer and loading buffer 6X (0.3 M Tris-HCl, 0.6 M DTT, 12% SDS, 0.06% Bromophenol blue, 50% Glycerol), boiled for 5 minutes to 95°C and subjected to electrophoresis using standard SDS-method. Proteins were transferred to a nitrocellulose membrane, stained with Ponceau S solution (Sigma-Aldrich #P7170) and blocked in PBS 0.1% Tween-20 buffer containing 10% of non-fat dried milk. After blocking, membranes were incubated overnight (O/N) at 4°C with primary antibodies prepared in PBS 0.1% Tween-20 buffer containing 2% of non-fat dried milk. Following day, membranes were washed with PBS 0.1% Tween-20 and incubated 1 hour at room temperature (RT) with peroxidase-conjugated secondary antibodies. Blots were developed using enhanced chemiluminescence reagent (Western Lighting Plus-ECL; Cytiva #28-9068-37), exposed to a radiograph film and develop using standard methods.

For immunohistochemistry, tissues were fixed in 10% buffered formalin (MilliporeSigma), except from brain which was fixed using 4%-PFA, and embedded in paraffin for routine histological analysis. IHC was performed on 2 μm paraffin sections by using an automated protocol developed for the DISCOVERYXT-automated slide-staining system (Ventana Medical Systems Inc.). All steps were performed in this staining platform by using validated reagents. Sections were stained with hematoxylin and eosin (H&E), or primary antibodies for specific markers (Suppl. Table 7). Appropriate biotinylated secondary antibodies were used to detect the primary antibodies, followed by incubation with streptavidin–horseradish peroxidase and diaminobenzidine system. Full slides were digitalized with a Zeiss AxioScan Z1 and analyzed by using the ZEISS Zen 2.3 Imaging Software (Zeiss) or QuPath-0.51 software.

For immunoprecipitation of liver samples, approximately 150-200 mg of tissue was processed. Tissue was placed in Precellys tubes containing zirconium beads and cold RIPA buffer (previously described). Then physical disaggregation of tissue was done using Precellys (Bertin Technologies). Following tissue disruption, samples were centrifuged at 13200 rpm for 15 minutes and 4°C to discard tissue leftovers and fat. Supernatant containing the protein extract was quantified and incubated with FLAG-conjugated magnetic beads (Merck, #M8823). 4 mg of protein was incubated O/N at 4°C with 50 μL of FLAG-conjugated magnetic beads using orbital shaker. Beads were then washed 10 times with 600 μL wash buffer (10 mM Tris-HCl pH 7.5, 0.5 mM EDTA, 150 mM NaCl). 80% of washed beads were collected and used for subsequent analysis by LC/MS. Remaining beads were eluted for WB. Elution by competition with FLAG peptide (3X) (Sigma Aldrich, # F4799) at 100 ng/μL diluted in TBS buffer was performed at RT for 30 minutes using an orbital shaker. Supernatant was collected, mix with LB 6X, boiled for 5 minutes and loaded for WB analysis.

For MEFs IP, 2×10^6^ cells were seeded in p150 plates. Following day, 1 μg/ml doxycycline was added to the medium and 24 hours later MEFs were collected as described before. 1mg of protein lysate was incubated with 30 μL FLAG-beads O/N. Following day, beads were washed, and elution was done using FLAG-peptide as previously described.

### Computational and statistical analysis

Preprocessing of IP-LC/MS and proteomics data was done using ProStaR software. Spectronaut (Biognosys) was used for phospho-proteomics data pre-processing. Bioinformatic analysis of gene set enrichment analysis (GSEA)(Fang *et al*, 2023), Metascape (Zhou *et al*, 2019), volcano plots and heatmaps were done using RStudio and Python 3.10. Phosphoproteomics analysis was performed using ProteomicsDB (Lautenbacher *et al*, 2022; Muller *et al*, 2025), Robust Inference of Kinase Activity (RoKAI App) (Yilmaz *et al*, 2021), Kinase Enrichment Analysis 3 (KEA3) (Kuleshov *et al*, 2021), Kinase Substrate Enrichment Analysis (KSEA)(Casado *et al*, 2013), and Posttranslational Modification Substrate Enrichment Analysis (PTM-SEA)(Krug *et al*, 2019). Statistical analysis was carried out using Prism 10 (GraphPad), python or RStudio using “rstatix” package. Statistical tests were performed using either two-tailed unpaired Student’s t-test, 1- or 2-way ANOVA (Dunnet’s, Tukey’s or Šidák’s multiple test) according to specifications in the figures. Data with p>0.05 were considered not statistically significant (ns); *, p<0.05; **, p<0.01; ***, p<0.001, ****, p<0.0001.

### Data accession

The mass spectrometry proteomics data have been deposited to the ProteomeXchange Consortium via the PRIDE partner repository with the dataset identifier PXD071218.

## Acknowledgements

We thank the Transgenic Unit of the CNIO for excellent work in the generation of the inducible mouse mode. L.R.L., and A.S.B. were supported by FPI contracts from the Spanish Ministry of Science and Innovation (MICINN PRE2019-088864). G.d.C was supported by grants from the AECC (LABAE16017DECA) and Ministry of Science and Innovation (MICINN /AEI/FEDER PID2021-125705OB-I00 and PID2024-159104OB-I00). M.M. lab was supported by research grants from Pfizer (Emerging Science Fund), MICINN/AEI/FEDER (PID2021-128726, PDC2022-133408-I00, RED2022-134792-T, RED2024-153635-T and PID2024-161681OB-100), MICINN-ISCIII (DTS24/00099), Foundation laCaixa (CaixaImpulse CI24-20693) and Comunidad de Madrid (Y2020/BIO-6519 and S2022/BMD-7437). This research was supported by AstraZeneca (JBID) and CIBER-Consorcio Centro de Investigación Biomédica en Red-(CIBERONC), Instituto de Salud Carlos III, MICINN. VHIO would like to acknowledge the Cellex Foundation for providing research facilities and equipment and the CERCA Program from the Generalitat de Catalunya for their support on this research. CNIO (CEX2019-000891-S) and VHIO (CEX2020-001024-S/AEI/10.13039/501100011033) are Centers of Excellence Severo Ochoa (Agencia Estatal de Investigación, MICINN).

## References

Acebron SP, Karaulanov E, Berger BS, Huang YL, Niehrs C (2014) Mitotic wnt signaling promotes protein stabilization and regulates cell size. Mol Cell 54: 663–674

An W, Zhang Z, Zeng L, Yang Y, Zhu X, Wu J (2015) Cyclin Y Is Involved in the Regulation of Adipogenesis and Lipid Production. PLoS One 10: e0132721

Bahn YJ, Yadav H, Piaggi P, Abel BS, Gavrilova O, Springer DA, Papazoglou I, Zerfas PM, Skarulis MC, McPherron AC et al (2023) CDK4-E2F3 signals enhance oxidative skeletal muscle fiber numbers and function to affect myogenesis and metabolism. J Clin Invest 133

Bakiri L, Hamacher R, Graña O, Guío-Carrión A, Campos-Olivas R, Martinez L, Dienes HP, Thomsen MK, Hasenfuss SC, Wagner EF (2017) Liver carcinogenesis by FOS-dependent inflammation and cholesterol dysregulation. J Exp Med 214: 1387–1409

Barron L, Courtney C, Bao J, Onufer E, Panni RZ, Aladegbami B, Warner BW (2018) Intestinal resection-associated metabolic syndrome. J Pediatr Surg 53: 1142–1147

Beard C, Hochedlinger K, Plath K, Wutz A, Jaenisch R (2006) Efficient method to generate single-copy transgenic mice by site-specific integration in embryonic stem cells. Genesis 44: 23–28

Beckonert O, Keun HC, Ebbels TM, Bundy J, Holmes E, Lindon JC, Nicholson JK (2007) Metabolic profiling, metabolomic and metabonomic procedures for NMR spectroscopy of urine, plasma, serum and tissue extracts. Nat Protoc 2: 2692–2703

Bø TH, Dysvik B, Jonassen I (2004) LSimpute: accurate estimation of missing values in microarray data with least squares methods. Nucleic Acids Res 32: e34

Casado P, Rodriguez-Prados JC, Cosulich SC, Guichard S, Vanhaesebroeck B, Joel S, Cutillas PR (2013) Kinase-substrate enrichment analysis provides insights into the heterogeneity of signaling pathway activation in leukemia cells. Sci Signal 6: rs6

Chen H, Jin C, Xie L, Wu J (2024) Succinate as a signaling molecule in the mediation of liver diseases. Biochim Biophys Acta Mol Basis Dis 1870: 166935

Chen L, Wang X, Cheng H, Zhang W, Liu Y, Zeng W, Yu L, Huang C, Liu G (2020) Cyclin Y binds and activates CDK4 to promote the G1/S phase transition in hepatocellular carcinoma cells via Rb signaling. Biochem Biophys Res Commun 533: 1162–1169

Da Silva F, Zhang K, Pinson A, Fatti E, Wilsch-Bräuninger M, Herbst J, Vidal V, Schedl A, Huttner WB, Niehrs C (2021) Mitotic WNT signalling orchestrates neurogenesis in the developing neocortex. Embo j 40: e108041

Davidson G, Niehrs C (2010) Emerging links between CDK cell cycle regulators and Wnt signaling. Trends Cell Biol 20: 453–460

Davidson G, Shen J, Huang YL, Su Y, Karaulanov E, Bartscherer K, Hassler C, Stannek P, Boutros M, Niehrs C (2009) Cell cycle control of wnt receptor activation. Dev Cell 17: 788–799

Delaglio F, Grzesiek S, Vuister GW, Zhu G, Pfeifer J, Bax A (1995) NMRPipe: a multidimensional spectral processing system based on UNIX pipes. J Biomol NMR 6: 277–293

Dohmen M, Krieg S, Agalaridis G, Zhu X, Shehata SN, Pfeiffenberger E, Amelang J, Bütepage M, Buerova E, Pfaff CM et al (2020) AMPK-dependent activation of the Cyclin Y/CDK16 complex controls autophagy. Nat Commun 11: 1032

Fang Z, Liu X, Peltz G (2023) GSEApy: a comprehensive package for performing gene set enrichment analysis in Python. Bioinformatics 39

Gassaway BM, Cardone RL, Padyana AK, Petersen MC, Judd ET, Hayes S, Tong S, Barber KW, Apostolidi M, Abulizi A et al (2019) Distinct Hepatic PKA and CDK Signaling Pathways Control Activity-Independent Pyruvate Kinase Phosphorylation and Hepatic Glucose Production. Cell Rep 29: 3394–3404.e3399

Gotoh S, Negishi M (2015) Statin-activated nuclear receptor PXR promotes SGK2 dephosphorylation by scaffolding PP2C to induce hepatic gluconeogenesis. Sci Rep 5: 14076

Holeček M (2023) Roles of malate and aspartate in gluconeogenesis in various physiological and pathological states. Metabolism 145: 155614

Hou S, Yu H, Liu C, Johnson AMF, Liu X, Jiang Q, Zhao Q, Kong L, Wan Y, Xing X et al (2024) Intestinal epithelial cell NCoR deficiency ameliorates obesity and metabolic syndrome. Acta Pharm Sin B 14: 5267–5285

Hughes CS, Moggridge S, Müller T, Sorensen PH, Morin GB, Krĳgsveld J (2019) Single-pot, solid-phase-enhanced sample preparation for proteomics experiments. Nat Protoc 14: 68–85

Jeong W, Jho EH (2021) Regulation of the Low-Density Lipoprotein Receptor-Related Protein LRP6 and Its Association With Disease: Wnt/beta-Catenin Signaling and Beyond. Front Cell Dev Biol 9: 714330

Jiang M, Gao Y, Yang T, Zhu X, Chen J (2009) Cyclin Y, a novel membrane-associated cyclin, interacts with PFTK1. FEBS Lett 583: 2171–2178

Johnson BA (2004) Using NMRView to visualize and analyze the NMR spectra of macromolecules. Methods Mol Biol 278: 313–352

Krug K, Mertins P, Zhang B, Hornbeck P, Raju R, Ahmad R, Szucs M, Mundt F, Forestier D, Jane-Valbuena J et al (2019) A Curated Resource for Phosphosite-specific Signature Analysis. Mol Cell Proteomics 18: 576–593

Kuleshov MV, Xie Z, London ABK, Yang J, Evangelista JE, Lachmann A, Shu I, Torre D, Ma’ayan A (2021) KEA3: improved kinase enrichment analysis via data integration. Nucleic Acids Res 49: W304–W316

Lautenbacher L, Samaras P, Muller J, Grafberger A, Shraideh M, Rank J, Fuchs ST, Schmidt TK, The M, Dallago C et al (2022) ProteomicsDB: toward a FAIR open-source resource for life-science research. Nucleic Acids Res 50: D1541–D1552

Lee Y, Dominy JE, Choi YJ, Jurczak M, Tolliday N, Camporez JP, Chim H, Lim J-H, Ruan H-B, Yang X et al (2014) Cyclin D1–Cdk4 controls glucose metabolism independently of cell cycle progression. Nature 510: 547–551

Lian H, Xu K, Chang A, Wang Y, Ma S, Cheng L, Zhao W, Xia C, Wang L, Yu G (2024) Loss of PTPN21 disrupted mitochondrial metabolic homeostasis and aggravated experimental pulmonary fibrosis. Respir Res 25: 426

Liu H, Shi H, Fan Q, Sun X (2016) Cyclin Y regulates the proliferation, migration, and invasion of ovarian cancer cells via Wnt signaling pathway. Tumour Biol 37: 10161–10175

Liu Z, Xue H, Wang Z, Zhao Y, Xu S, Dong X (2025) Cyclin Y interacts with Chk1 to activate RRM2/STAT3 signaling and promotes radioresistance in non-small cell lung cancer. Int J Biol Sci 21: 1999–2011

Luo H, Wang Q, Yang F, Liu R, Gao Q, Cheng B, Lin X, Huang L, Chen C, Xiang J et al (2023) Signaling metabolite succinylacetone activates HIF-1α and promotes angiogenesis in GSTZ1-deficient hepatocellular carcinoma. JCI Insight 8

Malumbres M (2014) Cyclin-dependent kinases. Genome Biol 15: 122

Martinez-Alonso D, Malumbres M (2020) Mammalian cell cycle cyclins. Semin Cell Dev Biol 107: 28–35

Mikolcevic P, Rainer J, Geley S (2012a) Orphan kinases turn eccentric: a new class of cyclin Y-activated, membrane-targeted CDKs. Cell Cycle 11: 3758–3768

Mikolcevic P, Sigl R, Rauch V, Hess MW, Pfaller K, Barisic M, Pelliniemi LJ, Boesl M, Geley S (2012b) Cyclin-dependent kinase 16/PCTAIRE kinase 1 is activated by cyclin Y and is essential for spermatogenesis. Mol Cell Biol 32: 868–879

Muller J, Bayer FP, Wilhelm M, Schuh MG, Kuster B, The M (2025) PTMNavigator: interactive visualization of differentially regulated post-translational modifications in cellular signaling pathways. Nat Commun 16: 510

Niehrs C, Acebron SP (2012) Mitotic and mitogenic Wnt signalling. EMBO J 31: 2705–2713

Petruzzelli M, Schweiger M, Schreiber R, Campos-Olivas R, Tsoli M, Allen J, Swarbrick M, Rose-John S, Rincon M, Robertson G et al (2014) A switch from white to brown fat increases energy expenditure in cancer-associated cachexia. Cell Metab 20: 433–447

Quandt E, Ribeiro MPC, Clotet J (2020) Atypical cyclins: the extended family portrait. Cell Mol Life Sci 77: 231–242

Saborano R, Eraslan Z, Roberts J, Khanim FL, Lalor PF, Reed MAC, Günther UL (2019) A framework for tracer-based metabolism in mammalian cells by NMR. Scientific Reports 9: 2520

Salazar-Roa M, Malumbres M (2017) Fueling the Cell Division Cycle. Trends in Cell Biology 27: 69–81

Schleucher J, Schwendinger M, Sattler M, Schmidt P, Schedletzky O, Glaser SJ, Sørensen OW, Griesinger C (1994) A general enhancement scheme in heteronuclear multidimensional NMR employing pulsed field gradients. J Biomol NMR 4: 301–306

Seo J, Hwang H, Choi Y, Jung S, Hong JH, Yoon BJ, Rhim H, Park M (2022) Myristoylation-dependent palmitoylation of cyclin Y modulates long-term potentiation and spatial learning. Prog Neurobiol 218: 102349

Shehata SN, Deak M, Morrice NA, Ohta E, Hunter RW, Kalscheuer VM, Sakamoto K (2015) Cyclin Y phosphorylation- and 14-3-3-binding-dependent activation of PCTAIRE-1/CDK16. Biochem J 469: 409–420

Shehata SN, Hunter RW, Ohta E, Peggie MW, Lou HJ, Sicheri F, Zeqiraj E, Turk BE, Sakamoto K (2012) Analysis of substrate specificity and cyclin Y binding of PCTAIRE-1 kinase. Cell Signal 24: 2085–2094

Shi K, Ru Q, Zhang C, Huang J (2018) Cyclin Y Modulates the Proliferation, Invasion, and Metastasis of Hepatocellular Carcinoma Cells. Med Sci Monit 24: 1642–1653

Stojanović O, Altirriba J, Rigo D, Spiljar M, Evrard E, Roska B, Fabbiano S, Zamboni N, Maechler P, Rohner-Jeanrenaud F et al (2021) Dietary excess regulates absorption and surface of gut epithelium through intestinal PPARα. Nature Communications 12: 7031

Sun T, Co NN, Wong N (2014) PFTK1 interacts with cyclin Y to activate non-canonical Wnt signaling in hepatocellular carcinoma. Biochem Biophys Res Commun 449: 163–168

Wang H, Nicolay BN, Chick JM, Gao X, Geng Y, Ren H, Gao H, Yang G, Williams JA, Suski JM et al (2017) The metabolic function of cyclin D3-CDK6 kinase in cancer cell survival. Nature 546: 426–430

Wieczorek S, Combes F, Lazar C, Giai Gianetto Q, Gatto L, Dorffer A, Hesse AM, Couté Y, Ferro M, Bruley C et al (2017) DAPAR & ProStaR: software to perform statistical analyses in quantitative discovery proteomics. Bioinformatics 33: 135–136

Willker W, Leibfritz, D., Kerssebaum, R. and Bermel, W. (1993) Gradient selection in inverse heteronuclear correlation spectroscopy. Magn Reson Chem 31: 287–292

Xue C, Chu Q, Shi Q, Zeng Y, Lu J, Li L (2025) Wnt signaling pathways in biology and disease: mechanisms and therapeutic advances. Signal Transduct Target Ther 10: 106

Yan F, Wang X, Zhu M, Hu X (2016) RNAi-mediated downregulation of cyclin Y to attenuate human breast cancer cell growth. Oncol Rep 36: 2793–2799

Yilmaz S, Ayati M, Schlatzer D, Cicek AE, Chance MR, Koyuturk M (2021) Robust inference of kinase activity using functional networks. Nat Commun 12: 1177

Yue W, Zhao X, Zhang L, Xu S, Liu Z, Ma L, Jia W, Qian Z, Zhang C, Wang Y et al (2011) Cell cycle protein cyclin Y is associated with human non-small-cell lung cancer proliferation and tumorigenesis. Clin Lung Cancer 12: 43–50

Zeng L, Cai C, Li S, Wang W, Li Y, Chen J, Zhu X, Zeng YA (2016) Essential Roles of Cyclin Y-Like 1 and Cyclin Y in Dividing Wnt-Responsive Mammary Stem/Progenitor Cells. PLoS Genet 12: e1006055

Zhang C, Li H, Han R (2020a) An open-source video tracking system for mouse locomotor activity analysis. BMC Res Notes 13: 48

Zhang L, Li X, Zhang N, Yang X, Hou T, Fu W, Yuan F, Wang L, Wen H, Tian Y et al (2020b) WDFY2 Potentiates Hepatic Insulin Sensitivity and Controls Endosomal Localization of the Insulin Receptor and IRS1/2. Diabetes 69: 1887–1902

Zhao X, Jiang M, Teng Y, Li J, Li Z, Hao W, Zhao H, Yin C, Yue W (2021) Cytoplasmic Localization Isoform of Cyclin Y Enhanced the Metastatic Ability of Lung Cancer via Regulating Tropomyosin 4. Front Cell Dev Biol 9: 684819

Zhou S, Zhang L, You Y, Yu K, Tie X, Gao Y, Chen Y, Yao F, Zhang R, Hao X et al (2025) eIF3f promotes tumour malignancy by remodelling fatty acid biosynthesis in hepatocellular carcinoma. J Hepatol

Zhou Y, Zhou B, Pache L, Chang M, Khodabakhshi AH, Tanaseichuk O, Benner C, Chanda SK (2019) Metascape provides a biologist-oriented resource for the analysis of systems-level datasets. Nat Commun 10: 1523

