## Supplementary Figures and Tables for "Cyclin Y Overexpression Drives a Fatal Metabolic Syndrome via Defective Glucose Homeostasis"

By Luis R. López-Sainz et al.

#### List of Supplementary Items

##### Supplementary Figures

Suppl. Fig. 1. Main alterations in a Cyclin Y over-expression mouse model.

Suppl. Fig. 2. Analysis of blood markers in CCNY-overexpressing mice.

Suppl. Fig. 3. CCNY induction affected other metabolites apart from glucose.

Suppl. Fig. 4. Metabolic alterations in CCNY-overexpressing MEFs.

Suppl. Fig. 5. Proteomic and biochemical analysis of CCNY-FLAG pull-downs.

Suppl. Fig. 6. Protein level changes in CCNY-overexpressing livers or liver tumors.

Suppl. Fig. 7. Phospho-proteomic changes in CCNY-overexpressing livers.

##### Supplementary Tables

Suppl. Table 1. Mass spectrometry analysis of proteins detected in CCNY-FLAG liver lysates from *Col1A1*(Y/Y) mice.

Suppl. Table 2. Changes in total protein levels in *Col1A1*(Y/Y) livers.

Suppl. Table 3. Quantification of changes in phospho-residues in *Col1A1*(Y/Y) livers.

Suppl. Table 4. Kinase pathways enriched as possible activities responsible for the changes in phosphoresidues observed in *Col1A1*(Y/Y) livers.

Suppl. Table 5. Pathways differentially represented in the analysis of phosphoresidues in *Col1A1*(Y/Y) livers after gene-centric redundant enrichment analysis using the Molecular Signatures Database (MSigDB)

Suppl. Table 6. Oligonucleotides used for genotyping of CCNY knock-in mice.

Suppl. Table 7. Antibodies used in this work.

### Supplementary Figures

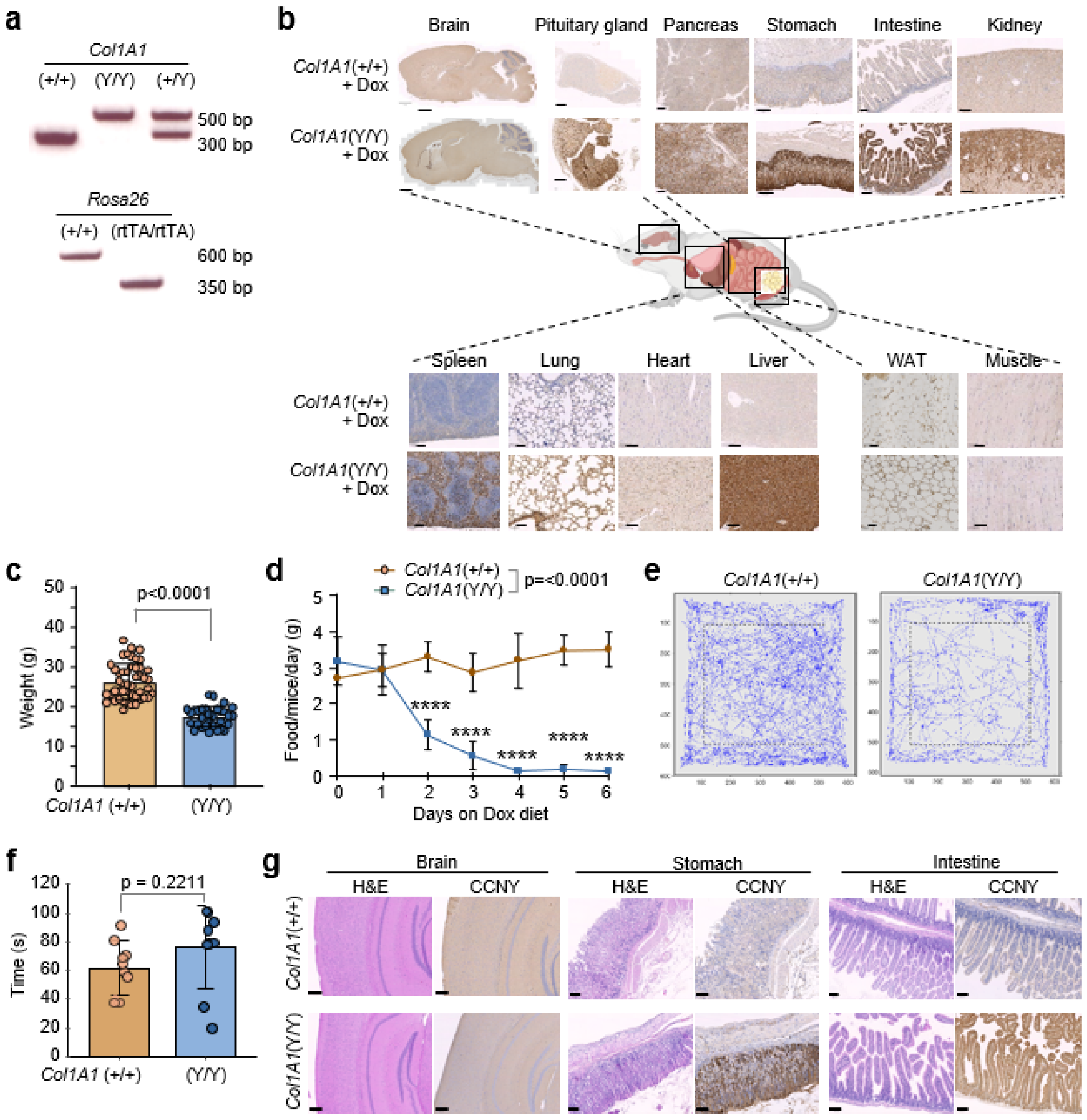


**Suppl. Fig. 1. Main alterations in a Cyclin Y over-expression mouse model. a)** Representative PCR products of the knockin alleles represented in Fig. 1a. **b)** Immunodetection of CCNY in histological sections of Dox-treated *Col1A1*(+/+) and *Col1A1*(Y/Y) mice in the indicated tissues. Scale bar, 1mm (brain); 250 µm (pituitary gland); 200 µm (kidney, stomach); 100 µm (heart, intestine, liver, muscle, pancreas, spleen, WAT); 50 µm (lung). **c)** Body weight of *Col1A1*(Y/Y) mice at human-end point (HEP) and in paired controls following Dox treatment (n=31-47 per group; two-tailed unpaired Student's t-test). **d)** Time course of daily food intake in grams after Dox diet (n= 8-9 per group). Data were analyzed using mixed-effect analysis with Šidák´s multiple comparison test (****p<0.0001) **e)** Representative trajectory plot during 15 minutes of control and *Col1A1*(Y/Y) mice (n= 10 mice per group). **f)** Rotarod performance of *Col1A1*(Y/Y) mice and paired controls. Time in seconds (s) to fall was plotted. (n = 10 per group). Data was analyzed using two-tailed unpaired Student's t-test**. g)** Representative hematoxylin & eosin (H&E) staining and immunodetection of CCNY in the indicated tissues and mouse models. In c,d,f data are expressed as mean ± SD

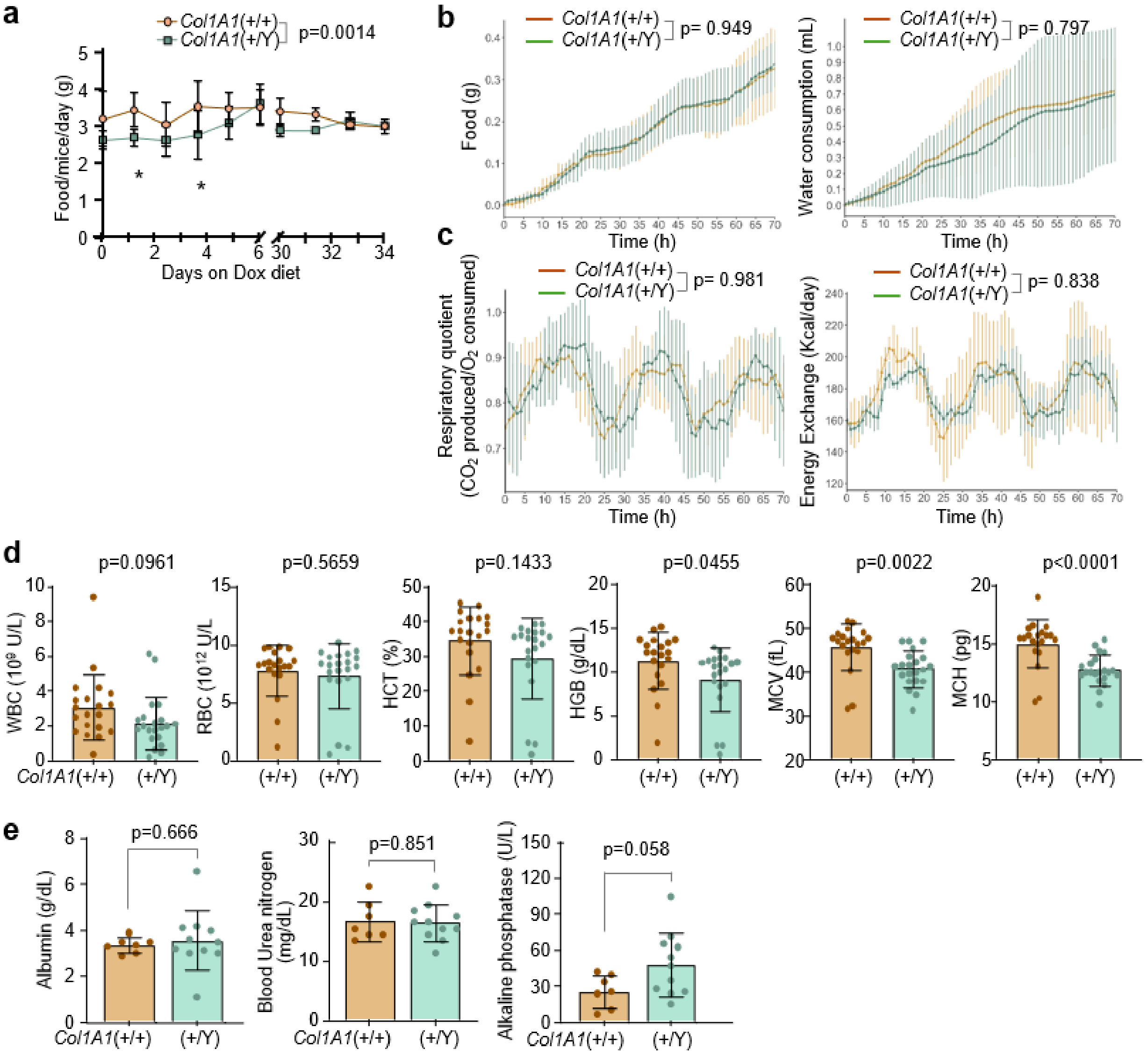


**Suppl. Fig. 2. Analysis of blood markers in CCNY-overexpressing mice. a)** Time course of blood glucose levels following Dox diet administration (n=5-8 mice per genotype). Data were analyzed using mixed-effect analysis with Šidák´s multiple comparison test (*p<0.05). **b)** Time course analysis of food and water consumption in control and *Col1A1*(+/Y) mice using metabolic cages. **c)** Time course analysis of respiratory quotient and energy exchange in control and *Col1A1*(+/Y) mice using metabolic cages. In b,c, mice were on Dox diet for 100 days (n = 3 per genotype), and each time point was analyzed using two-tailed unpaired t-test and mixed-effect analysis with Šidák´s multiple comparison test for condition effect analysis. **d)** Blood analysis of the indicated parameters. Every dot represents a different mouse. Data were analyzed using unpaired Student's t-test. **e)** Bloodstream levels of albumin, blood urea nitrogen, and alkaline phosphatase in *Col1A1*(+/Y) mice at humane endpoint and paired controls. Each dot represents a single mouse (n = 5-11 mice per group). Data are expressed as means ± SD, and were analyzed using two-tailed unpaired Student's t-test.

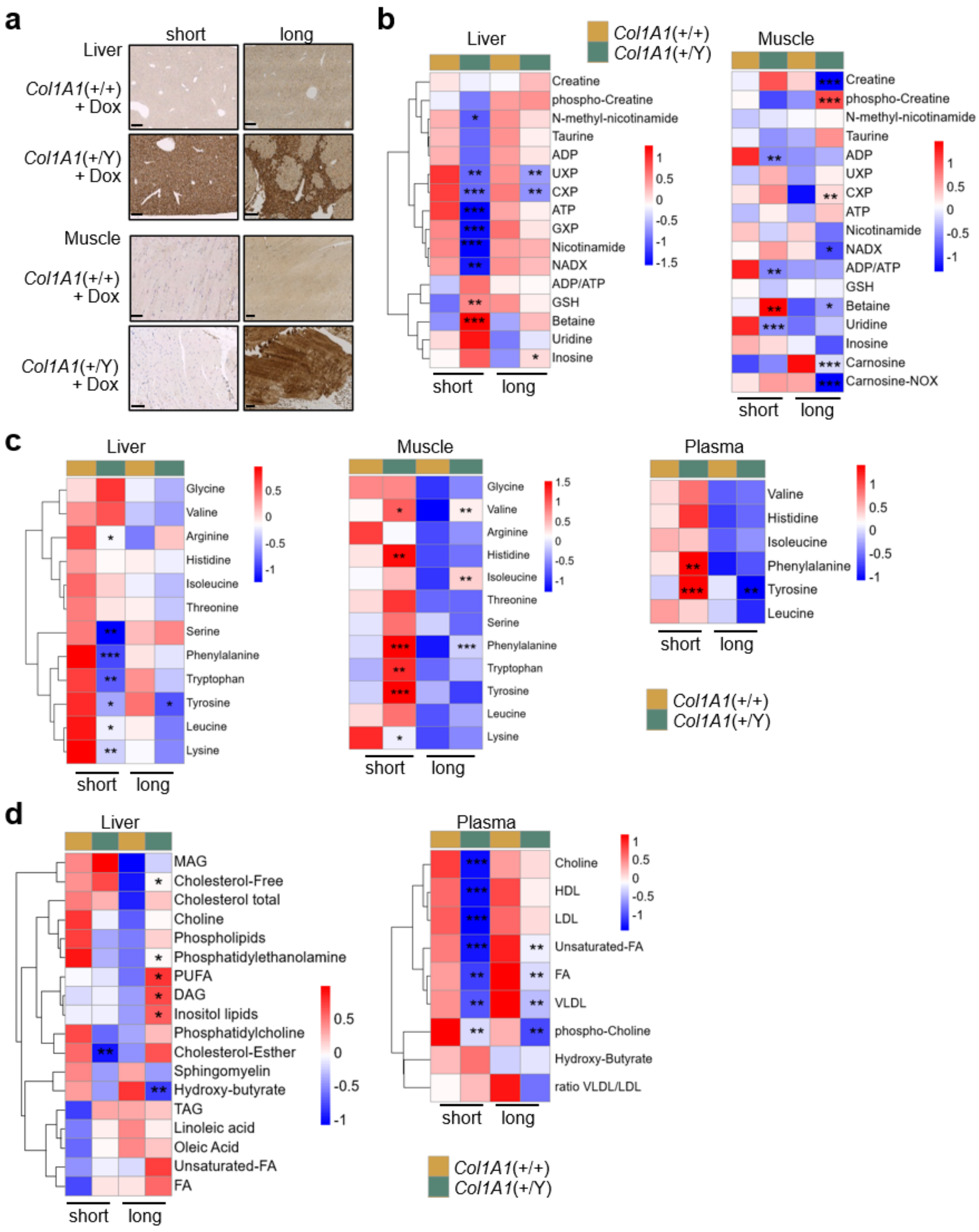


**Suppl. Fig. 3. CCNY induction affected other metabolites apart from glucose. a)** Immunodetection of CCNY in liver and muscle histologies from *Col1A1*(+/Y) mice and their matched controls, after short of long exposure to Dox as described in Fig. 3f. Scale bar, 200 µm (liver); 100 µm (muscle). **b-d)** Heatmap of z-score corrected data from NMR measurements in the indicated samples of molecules related to nucleotide metabolism (**b**), amino acids (**c**) or lipids (**d**). Groups and time points are annotated (n=5-6 mice per group). Statistical comparisons were analyzed using two-tailed unpaired Student's t-test of CCNY-overexpressing versus control mice subjected to short or long Dox exposure. *p<0.05; **p<0.005; ***p<0.0005.

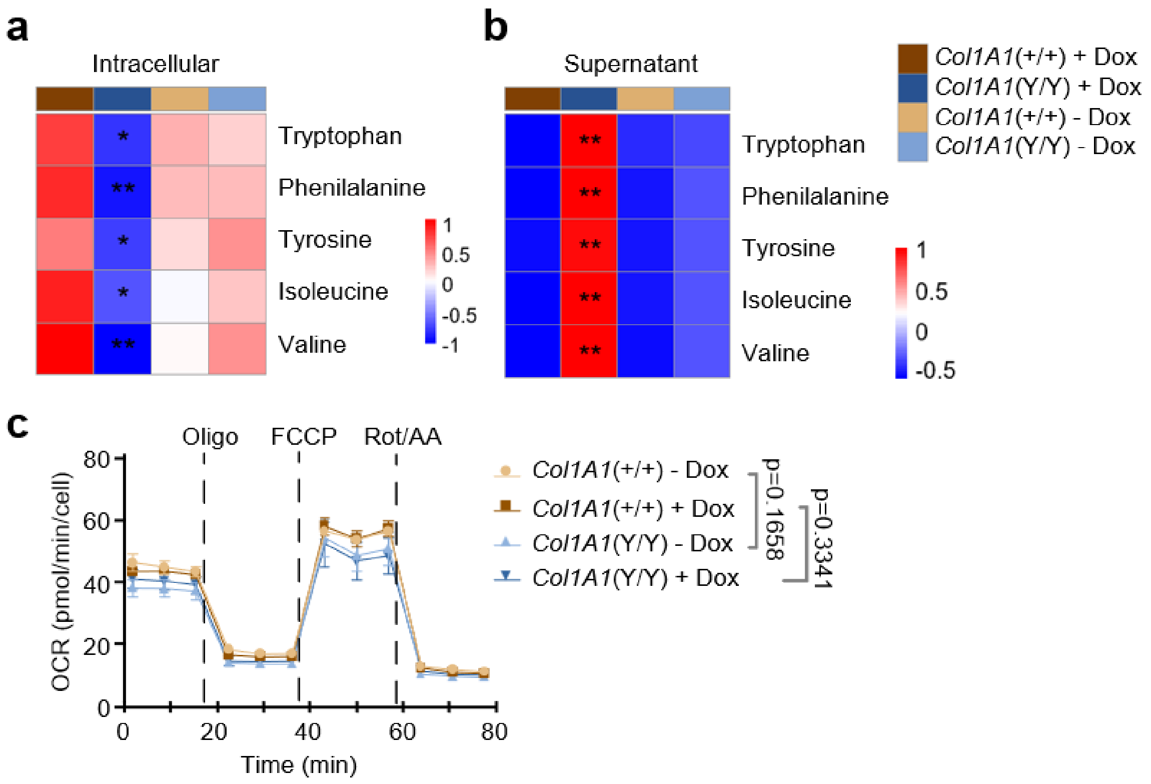


**Suppl. Fig. 4. Metabolic alterations in CCNY-overexpressing MEFs.** (**a,b)** Heatmap of z-score normalized data from NMR measured amino acid concentrations in mouse embryonic fibroblasts (MEFs) treated with or without Dox. Panel (**a**) represents intracellular amino acids and (**b**) the supernatant (n=4-6 cultures per group). Data were analyzed using two-tailed unpaired Student's t-test of CCNY-overexpressing vs. control MEFs; *p<0.05; **p<0.005. **c)** Seahorse analysis of *Col1A1*(+/+) and *Col1A1*(Y/Y) MEFs treated with or without Dox. Time course analysis of oxygen consumption rate (OCR) with the specific inhibitors administrated in each experiment (n= 4-6 cultures per group). Data is expressed as means ± SEM and statistics were analyzed with two-way ANOVA with Šidák´s multiple comparison test of compared conditions.

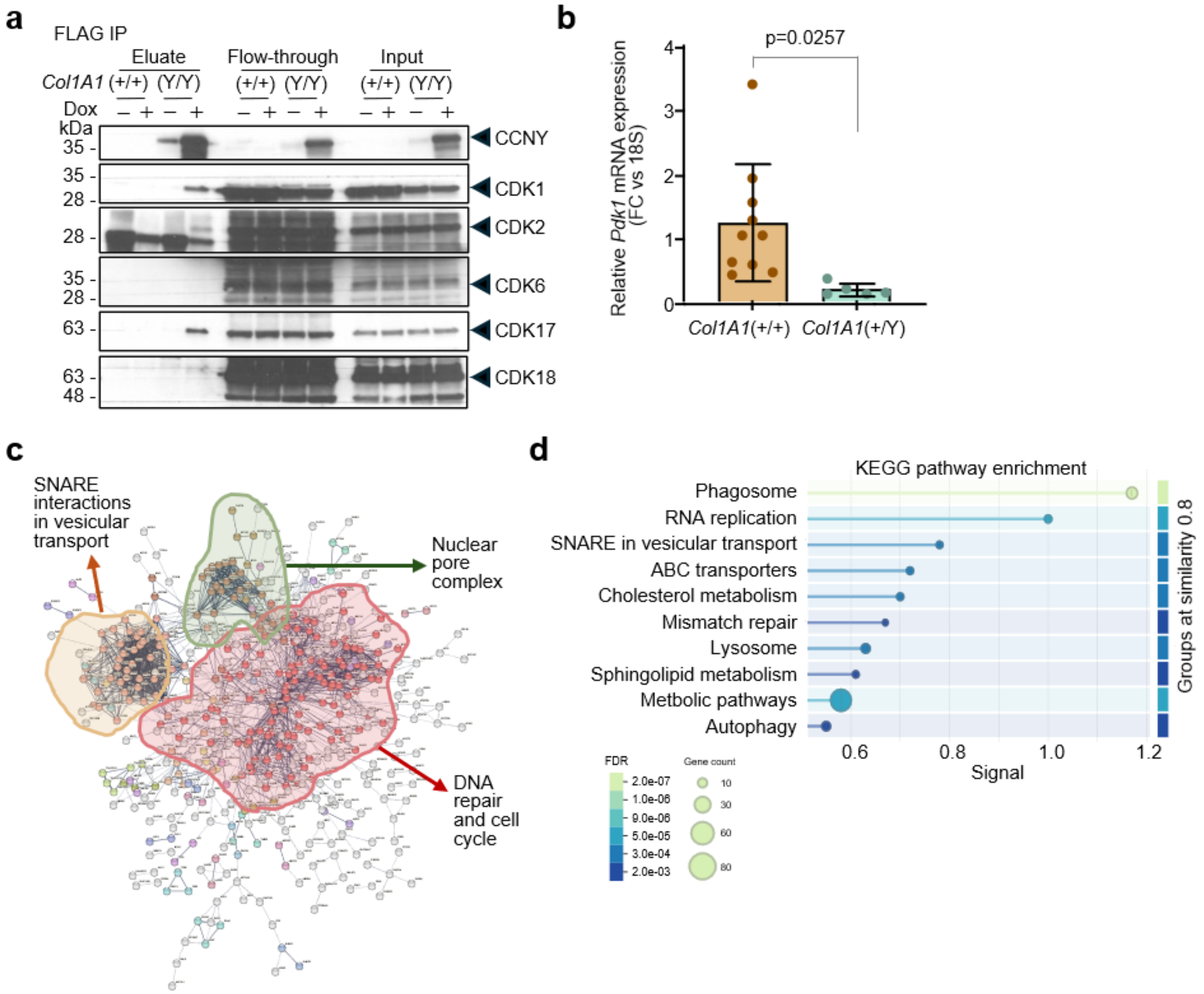


**Suppl. Fig. 5. Proteomic and biochemical analysis of CCNY-FLAG pull-downs. a)** FLAG pulldown of *Col1A1*(+/+) and *Col1A1*(Y/Y) lysates from mouse embryonic fibroblasts (MEFs) treated for 24 hours with/without Dox. CCNY and CDKs were immunoblotted in eluate, flow-through and input fractions. Black rows indicate the specific bands corresponding to each protein. **b)** Relative *Pdk1* mRNA levels in livers from control and *Col1A1*(+/Y) mice normalized versus the levels of 18S ribosomal gene. Fold change is represented. Each dot represents a single mouse. Data are expressed as means ± SD and were analyzed using two-tailed unpaired Student's t-test. **c)** Interacting network generated in STRING database from significant over-represented proteins in Fig. 5b. DBSCAN clustering from STRING shows the three main annotated clusters. **d)** Functional enrichment visualization from STRING database of KEGG pathways enriched in proteins detected after CCNY-FLAG pull-downs in liver samples from CCNY-overexpressing mice. Signal, false discovery rate (FDR) and gene count are annotated for each pathway.

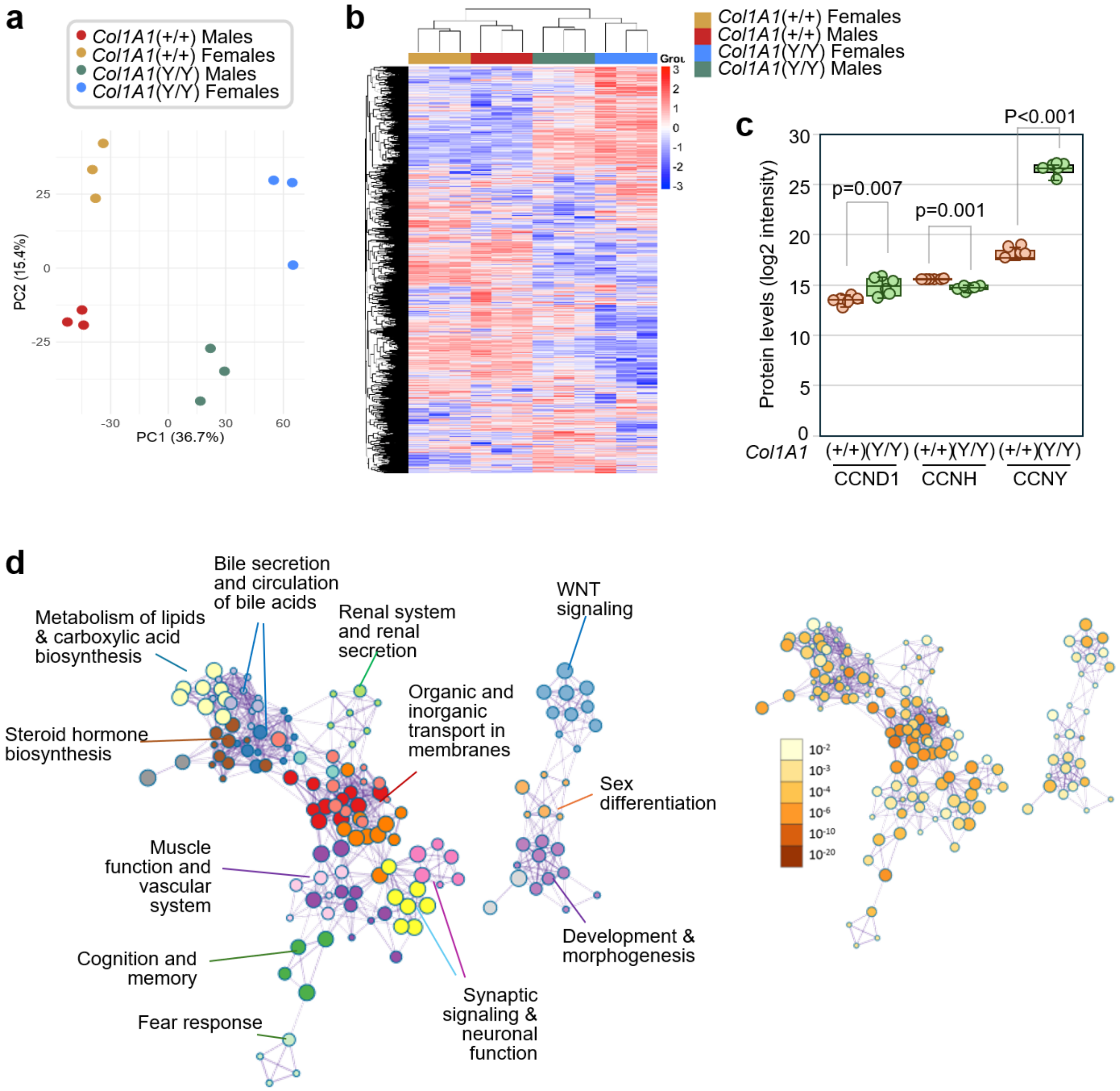


**Suppl. Fig. 6. Protein level changes in CCNY-overexpressing livers or liver tumors. a)** Principal component analysis (PCA) of z-score corrected protein levels from three mice per genotype and gender. PC1 and PC2 variances are represented in percentage. **b)** Clustered heatmap of z-score corrected data of protein levels (rows) in the indicated groups (columns). **c)** Quantitative comparison of peptide levels for the indicated cyclins. Every dot indicates a separate mouse, and data are represented as mean ± SD. Unpaired Student's t-test. **d)** Analysis of pathways significantly enriched in CCNY-overexpressing liver hepatocellular carcinoma compared with CCNY-low liver tumors. Selected names cluster individual pathways (dots) with proteins in common. The statistical analysis of these pathways is color-coded in the panel to the right (Metascape analysis). Data from TCGA LIHC (liver hepatocellular carcinoma) comparing tumors with quartil 3 (Q3) versus Q1 CCNY expression at the mRNA level.

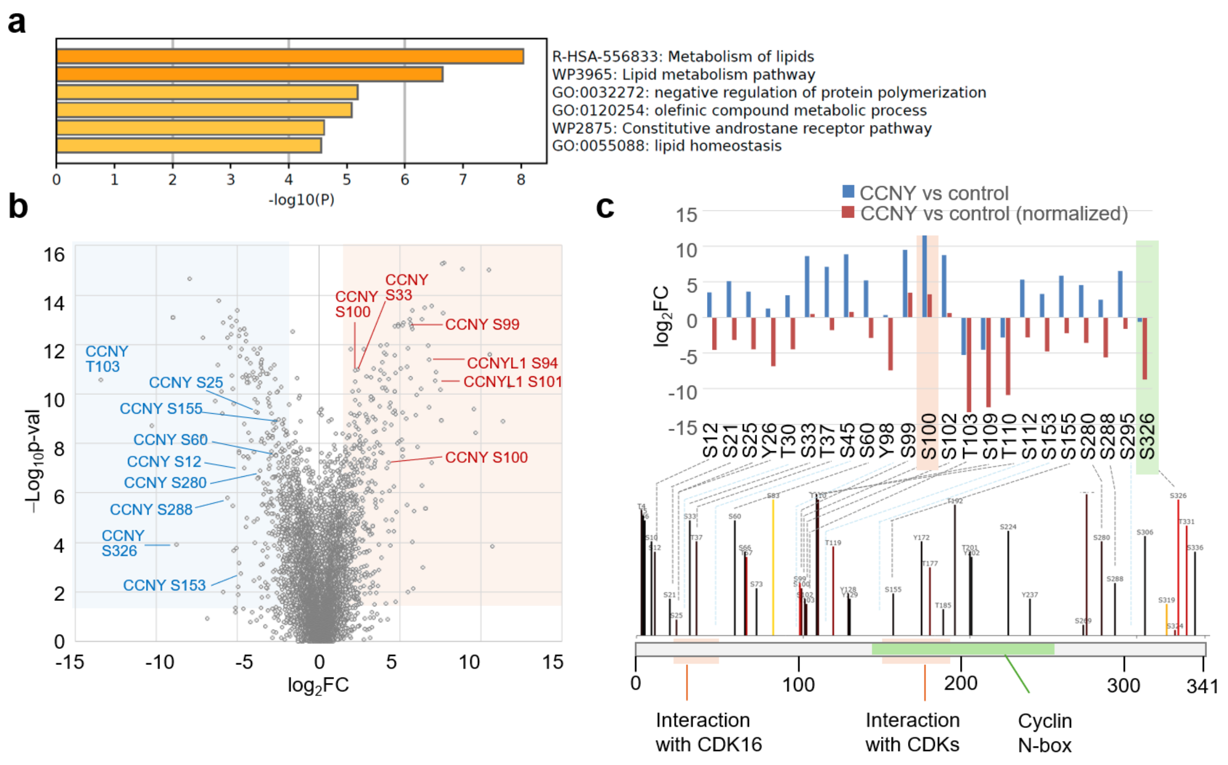


**Suppl. Fig. 7. Phospho-proteomic changes in CCNY-overexpressing livers. a)** Pathways preferentially enriched in proteins with the phosphosites significantly dysregulated when comparing female versus male *Col1A1*(Y/Y) livers. The plot shows the main over-represented pathways (Metascape analysis). **b**) Volcano plot of phosphosites upregulated in CCNY-overexpressing livers. Sites with logFC >2 or <–2 and p-value <0.05 were selected as upregulated (red) or downregulated (blue), respectively. CCNY and CCNYL1 up (red) or down (blue) phosphosites are annotated. **c**) Schematic representation of the main domains in the human CCNY protein with phosphoresidues represented in The Kinase Library (https://kinase-library.mit.edu/home; bottom). The upper panel shows the fold change before (blue) and after (red) normalization for changes in total protein levels in CCNY-overexpressing livers compared to controls. The blue dotted lines indicate phosphoresidues not represented in The Kinase Library. Residues S100 and S326 control CCNY function and are discussed in the main text.

#### Supplementary Tables

**Suppl. Table 1. Mass spectrometry analysis of proteins detected in CCNY-FLAG liver lysates from *Col1A1*(Y/Y) mice.**

**Suppl. Table 2. Changes in total protein levels in *Col1A1*(Y/Y) livers.**

**Suppl. Table 3. Quantification of changes in phospho-residues in *Col1A1*(Y/Y) livers.**

**Suppl. Table 4. Kinase pathways enriched as possible activities responsible for the changes in phosphoresidues observed in *Col1A1*(Y/Y) livers.**

**Suppl. Table 5. Pathways differentially represented in the analysis of phosphoresidues in *Col1A1*(Y/Y) livers after gene-centric redundant enrichment analysis using the Molecular Signatures Database (MSigDB)**

**Suppl. Table 6. Oligonucleotides used for genotyping of CCNY knock-in mice.**

| Allele | Primer sequence | Annealing Tº | Size band bp (wt/mut) |
|---|---|---|---|
| Col1A1 (CCNY) | GCACAGCATTGCGGACATGC | 60ºC | 300 bp/ 500 bp |
| | CCCTCCATGTGTGACCAAGG | | |
| | GCAGAAGCGCGGCCGTCTGG | | |

**Suppl. Table 7. Antibodies used in this work.**

| Antigen | Reference | Company | Dilution (WB/IP) | Dilution (IHC) |
|---|---|---|---|---|
| CCNY | 18042-1-AP | Proteintech | 1:1000 (WB/IP) | - |
| CCNY | orb1275400 | Biorbyt | - | 1:2000 |
| CDK1 | HPA003387 | Merck | 1:1000 (WB/IP) | - |
| CDK17 | PA5-99841 | Thermo Scientific | 1:500 (WB) | - |
| CDK18 | NBP1-92249 | Novus Biologicals | 1:500 (WB/IP) | - |
| CDK2 | sc-6248 | Santa Cruz | 1:1000 (IP) | - |
| CDK4 | 12790 | Cell Signaling | 1:1000 (WB/IP) | - |
| CDK6 | ab124821 | Abcam | 1:1000 (WB/IP) | - |
| FLAG | F1804 | Sigma Aldrich | - | 1:400 |
| GLUT1 | 21829-1-AP | Proteintech | 1:1000 (WB) | - |
| PDK1 | 3062T | Cell Signaling | 1:1000 (IP) | - |
| PDK4 | PA5-102685 | Thermo Scientific | 1:1000 (IP) | - |
| Vinculin | V9131-2ML | Sigma Aldrich | 1:20.000 (WB) | - |
